# Loss of PIF7 attenuates shade and elevated temperature responses throughout the lifecycle in pennycress

**DOI:** 10.1101/2025.03.17.643809

**Authors:** Vanessica Jawahir, Salma Adam, Marcus Griffiths, Tad Wesley, Win Phippen, Nicholas Heller, Bhabesh Borphukan, Karen Sanguinet, Dmitri A. Nusinow

## Abstract

Pennycress (*Thlaspi arvense*) is being developed as a winter annual intermediate oilseed bioenergy crop in the Midwest during typical fallow periods. Crucial work remains to domesticate and optimize pennycress for incorporation into cropping systems and increasing resilience to rising temperatures. We found that increased planting density reduces biomass and hastens time to flowering and maturity, which are associated with shade avoidance responses. In controlled conditions, we found that pennycress elongates in response to foliar shade and increased ambient temperatures (28 °C). We applied the knowledge base from *Arabidopsis thaliana* to manipulate genes in the PHYB signaling pathway to simultaneously decrease the shade avoidance response during interseeding and tissue responses to elevated temperatures. Evaluation of CRISPR alleles of PIF7 shows that *pif7* reduces organ elongation to competition and heat cues and retains a compact rosette when exposed to shade or elevated temperature and their combination. Crucially, yield and oil content were unaltered in *pif7* and plants maintained earlier flowering in stress conditions. Furthermore, indicators of plant health, such as hue, chlorophyll indices, and root system architecture, were improved between wild type and *pif7*. This is evidence that plant architecture and physiological health can be uncoupled under competition and heat conditions, supporting our efforts to attenuate morphological responses to environmental cues. We propose this strategy for reducing SAR, improving pennycress performance at high densities, for during interseeding establishment in standing crops, and in a warming climate.

## Introduction

Agriculture is tasked with increasing production without expanding land use to sustainably produce feed, fiber, and biofuel for the rapidly growing human population (Ranganathan et al. 2018). The required increases in yield can be achieved by improving crop genetics, increasing planting density, and planting more frequently. Recently, improved yield has come from selecting for tolerance to increased planting densities (Duvick 2005). However, planting densities must be carefully optimized as excessive density plantings trigger the shade avoidance response (SAR), a syndrome resulting from perceived competition for light. SAR, a major limitation to crop planting density, leads to elongated growth, altered tillering, increased lodging, and early flowering, and can result in the underperformance of the shorter species during intercropping (Lu et al. 2023; Li et al. 2021; Tang & Liesche 2017; Liu et al. 2021). Elongation and early flowering are evolutionarily advantageous to increase the fitness of individuals but are undesirable in crop monoculture systems as elongated plants are more prone to lodging, and allocation of resources to these competitive traits often reduces yield (Tang & Liesche 2017; Wille et al. 2017; Raza et al. 2020; Roig-Villanova & Martínez-García 2016; Schmitt et al. 1999).

The effects of SAR have been partially reduced in cereals by breeding for upright leaf angle high in the canopy to better capture light and reduce canopy shading (Wille et al. 2017; Cao et al. 2022). This altered plant architecture allows for significantly increased *Zea maize* planting density from 30,000 plants/hectare to 80,000 plants/hectare, a major factor in yield gains (Duvick 2005; Assefa et al. 2018). Improvements in leaf angle have slowed, suggesting this trait is at near peak optimization in maize (Elli et al. 2023), and increased tolerance to higher planting density will need to be accomplished through alternative mechanisms. Leaf angle is not a uniform target for trait improvement in all crops, especially rosette-forming crops. An alternative strategy is to attenuate SAR and diminish changes that occur in response to high planting density. Studies in *Arabidopsis thaliana* have shown that reduction of SAR improved monoculture performance at high densities and increased weed suppression (Pantazopoulou et al. 2021), supporting that SAR attenuation is a viable strategy.

The molecular mechanism of SAR has been well studied in Arabidopsis and is a rich resource to mine for candidates for targeted improvement. From the Arabidopsis literature, it is understood that plants continually monitor the environment for changes in light quality, quantity, and temperature through phytochromes to initiate transcriptional responses to respond to the environmental signals (Oravec & Greenham 2022; Jung et al. 2016). Signaling through the five phytochromes (phyA to E) in Arabidopsis regulates germination, chlorophyll biosynthesis, photosynthesis, circadian rhythms, shade avoidance, hypocotyl elongation, and flowering times (Sharrock & Quail 1989; Clack et al. 1994; Sharrock 2008; Kami et al. 2010). Phytochromes are converted to the Pfr (active) form upon red (R) (660 nm) light exposure and reverted to the Pr (inactive) form either upon far-red (Fr) (730 nm) light exposure, exposure to elevated temperatures (thermal reversion), or by incubation in the dark in a process termed dark reversion (Rockwell et al. 2006). Active levels of PHYB respond to the R:Fr ratio. R:Fr in sunlight is approximately 1.15 but is reduced in shade as green canopies are effective at absorbing R light but allow Fr to pass through (Franklin & Whitelam 2005; Smith 1982). Thus, a low R:Fr is the cue to an individual that it is being shaded by other plants and should initiate SAR to overtop neighboring plants.

Phytochromes also function as thermosensors due to temperature-dependent PHYB thermal reversion from Pfr to Pr, clarifying their role in plant responses to elevated temperatures (Legris et al. 2016; Jung et al. 2016). Arabidopsis grown in warm temperatures (28°C) display elongated hypocotyls, petioles, and roots, altered leaf angle, and flowers earlier compared to growth at lower temperatures (Delker et al. 2022; Casal & Balasubramanian 2019; Fernández et al. 2016). Thermomorphogenesis and SAR are phenotypically similar in Arabidopsis, due to PHYB’s shared perception of light quality and temperature and initiation of a common signaling cascade to regulate growth. Combining warm temperatures and shade has been reported to have synergistic effects on Arabidopsis hypocotyl elongation (Burko et al. 2022).

PHYB modulates plant growth through its interaction with PHYTOCHROME INTERACTING FACTORS (PIF) transcription factors to regulate their post-translational sequestration and turnover (Nozue et al. 2007; Jung et al. 2016; Bauer et al. 2004; Lorrain et al. 2008; Park et al. 2012; Park et al. 2018; Kim et al. 2024). PhyB in the active Pfr state binds and sequesters PIFs, preventing their ability to induce gene expression. Repression of PIF activity is relieved when the phytochromes are converted to Pr by either far-red light, darkness, or high temperatures, thereby stabilizing the PIFs and inducing elongation programs. The PIFs induce the expression of auxin biosynthesis genes, leading to organ elongation, which alters plant morphology in response to changing light and temperature conditions (Legris et al. 2019; Pham et al. 2018; Balcerowicz 2020; Li et al. 2012; Sessa et al. 2018). Individual PIFs have both distinct and redundant functions to modulate responses in specific environments, allowing for fine-tuned growth responses (Pham et al. 2018; Balcerowicz 2020). PIF7 participates in the signaling cascade to reprogram developmental responses to both shade and elevated temperatures. In Arabidopsis, *pif7* mutants have reduced hypocotyl elongation in shade, elevated temperatures, and in combination (Burko et al. 2022). Similarly, *Cardamine hirsuta,* a shade and heat-tolerant plant, does not elongate its hypocotyl when grown in shade and elevated temperatures. *C. histura*’s blunted elongation response is attributed to reduced PIF7 availability compared to the shade and temperature-sensitive Arabidopsis (Paulišić et al. 2021), supporting that modulation of PIF7 activity can confer tolerance to shade and elevated temperature. *pif7* mutants also reduce shade-induced leaf hyponasty, resulting in quicker canopy closure, and were superior to wild type at suppressing the growth of invading plants when grown in field-like conditions (Pantazopoulou et al. 2021).

We sought to test if we can simultaneously reduce SAR and thermomorphogenesis by mutating PIF7 in pennycress (*Thlaspi arvense* L.; field pennycress). The pennycress genome is small, diploid, and more than 70% of the predicted peptides of the pennycress genome have more than 80% sequence similarity to those of Arabidopsis (Dorn et al. 2015). Pennycress is in the early stages of domestication as an intermediate oilseed crop with the dual benefit of soil protection during the winter. The September through October planting window is a crucial predictor of yield. Therefore, in northern US regions, growers will likely need to interseed pennycress into standing crops or in crop residue, forcing pennycress seedlings to establish themselves under shaded conditions. Pennycress had superior stand establishment when interseeded into soy rather than maize, which was attributed to higher light quality and light penetration through the soy canopy (Mohammed et al. 2020). We hypothesize that reduced SAR in pennycress will allow for improved intercropping, tolerance to higher planting densities, and greater ground cover during the winter. Altering the pathways regulating SAR could also impact thermotolerance, given they share components, and altered lines could be more tolerant to the fluctuating temperatures in fall and spring. Pennycress is a prime species to test the benefits of reducing SAR since overcoming interseeding challenges is relevant to the crop’s success.

In this study, we found that high planting densities negatively impact pennycress biomass and that pennycress responds to shade and elevated temperature. We mutated PIF7 and showed that *pif7* has reduced organ elongation when exposed to shade or elevated temperature, and their combination. At high densities, *pif7* has reduced elongation, supporting that reducing SAR is a promising strategy for improving plant performance at high densities.

## Results

### Planting density affects whole plant morphology and flowering time, but not yield

Optimizing planting density for any crop is necessary to balance maximal yield potential with competition for nutrients and light. To explore effects of planting density on pennycress morphology and ground cover, ARV1*tt8*, an early commercial pennycress line, was sown in four replicate plots at 3, 5, 7, and 9 lbs per acre in Macomb, IL, in 2023. Morphology traits were monitored throughout the 2023-2024 field season. At maturity, five plants from the center of each plot were harvested to characterize morphological differences. Plants at 3 lbs/acre produced the most tillers and had the highest biomass (Figure 1). Plants sown at 5 lbs/acre or higher developed significantly fewer tillers and had lower biomass. Changes in morphology were also apparent at the rosette stage before plants had entered winter dormancy. Sixty-four days after planting, five rosettes from each plot were harvested and imaged to measure rosette traits and leaf number. Rosettes from each planting density had statistically the same leaf area, but became more elongated at 7 and 9 lbs/acre, reducing solidity (Supplementary Figure 1). A reduction in solidity indicates that there is more space between the leaves as the convex hull contains a lower percentage of plant material.

**Figure 1.**
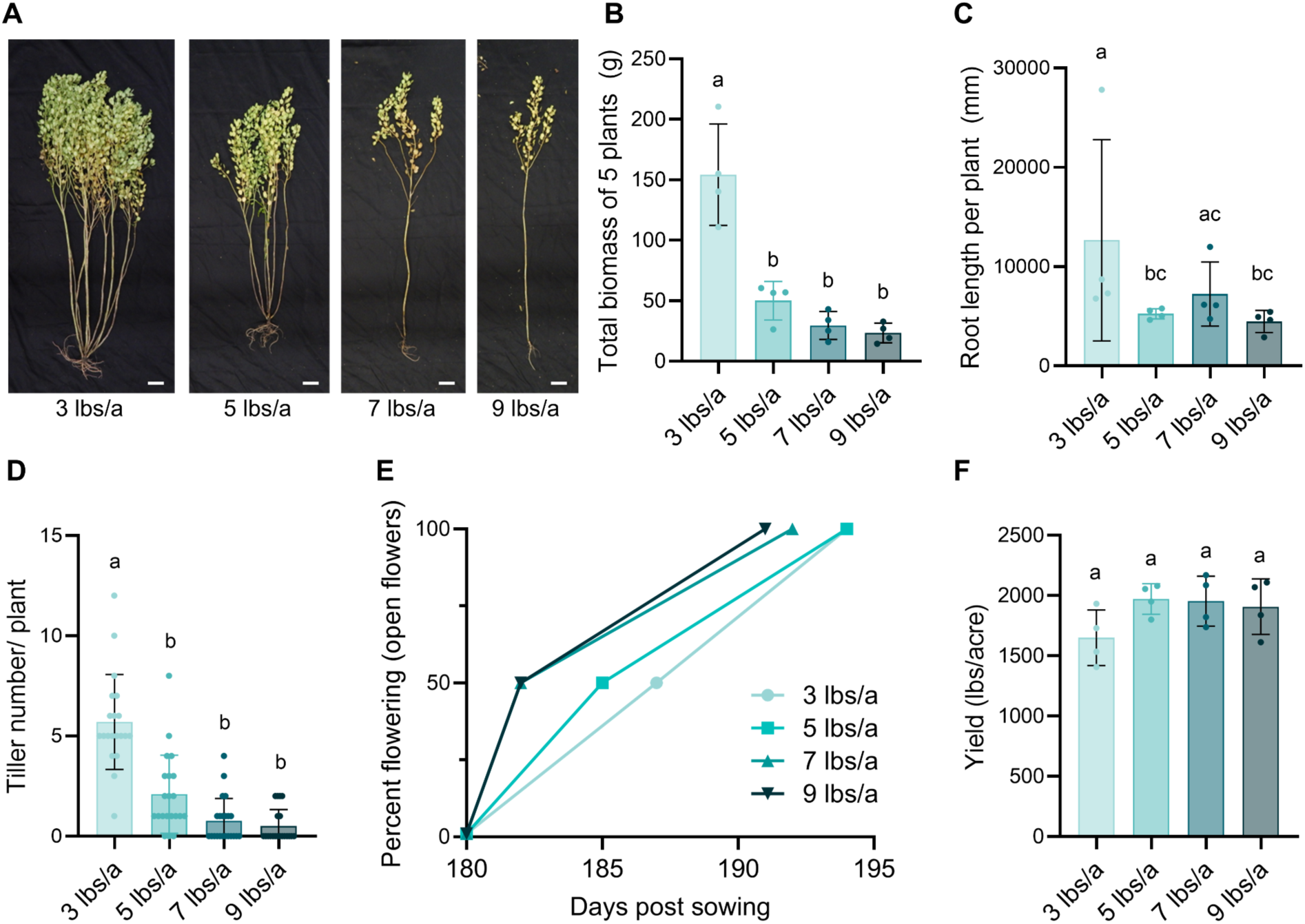
Biomass and yield negatively correlate with seeding density. **(A)** Four plots were sown at 3, 5, 7, and 9 lbs/a. At maturity, plants were harvested from the middle of the plot and imaged. Representative images of individuals from plots are shown. **(B)** Total dried biomass of 5 plants taken from each plot. Statistical significance determined by 2-tailed Welch’s t-test (n=4, p<0.05). **(C)** Total root length per plant was determined from soil cores from the middle of each plot. Rhizovision Explorer measured root length per core and divided by the number of tap roots present. Statistical significance determined by 2-tailed Mann-Whitney test (n=4, p<0.05) **(D)** The number of tillers per plant was counted from harvested individuals. Statistical significance determined by 2-tailed Mann-Whitney test. (n ≥ 20, p<0.05). **(E)** Plots were scored for the dates when each reached 50% and 100% flowering. **(F)** Plots were harvested, and the yield for each plot was determined. Statistical significance determined by 2-tailed Welch’s t-test (n=4, p<0.05). Common letters indicated no significant difference.

Changes in individual plant biomass were significant in both above- and below-ground biomass. Root cores were collected from the center of each plot 179 days post-sowing, and the number of plants per root core was determined by counting the number of tap roots present in each core. Plots of 5, 7, and 9 lbs/acre had the greatest number of plants per root core, indicating that higher-density plots had more plants. (Supplementary Figure 1). At 3 lbs/a, individual plants had longer total root length, larger average root diameter, and more biomass than plants at higher densities (Figure 1 and Supplementary Figure 1). Overall, we observed reduced root and shoot biomass per plant when grown at higher densities, demonstrating morphological changes secondary to increased planting densities.

Pennycress is a dual-purpose intermediate crop with large rosettes that can protect the soil, similar to cover crops. We assessed this secondary benefit of pennycress by determining the percent plant coverage of the total plot area. Drone images of the plots were collected every week starting in February after planting. Percent green coverage was determined by applying a color threshold to calculate the plot percentage covered by plant versus bare soil. The first drone images at day 149 post-sowing are representative of the percent plot protection during the majority of the winter months, with minimal above-ground growth during this time. There were no statistical differences in percent green coverage for all planting densities on day 149, and all time points after day 187 post-sowing (Supplementary Figure 1). The higher planting densities had greater overall percent green cover during the growth and bolting phase. Green cover plateaued after 200 days post-sowing and began to decrease starting day 235 post-planting, likely due to senescence at maturity.

Plots at all densities began to open flowers at day 180 post-sowing. Plots sown at 7 and 9 lbs/acre reached 50% and 100% flowering earlier than the 3 lbs/a plots. At maturity, all plots were harvested, and the average yield from all plot densities was statistically the same. This indicates that increasing pennycress planting density above 3 lbs/a does not increase yield per plot (Figure 1). We hypothesize that the morphological responses seen at higher densities reduce per-plant yield and indicate plant competition within the plot.

### Pennycress elongates in response to shade and elevated temperatures

The reduced tillering and early flowering phenotypes seen at high densities (Figure 1) are indicative of SAR and occur in the presence of canopy and neighbor shade in shade-intolerant plants (Wille et al. 2017; Franklin 2008; Cerdán & Chory 2003). Therefore, we hypothesized that pennycress is shade intolerant. Pennycress responses to shade were measured in the MN106 reference background as this line has a sequenced genome (Dorn et al. 2015; Nunn et al. 2022). Hypocotyl elongation was measured for seedlings germinated in white light, then supplemented with far-red light to reduce the Red:Far red (R:FR) from 2.1 to 0.2 during the day. Seedlings increased hypocotyl length under shade (Figure 2).

**Figure 2.**
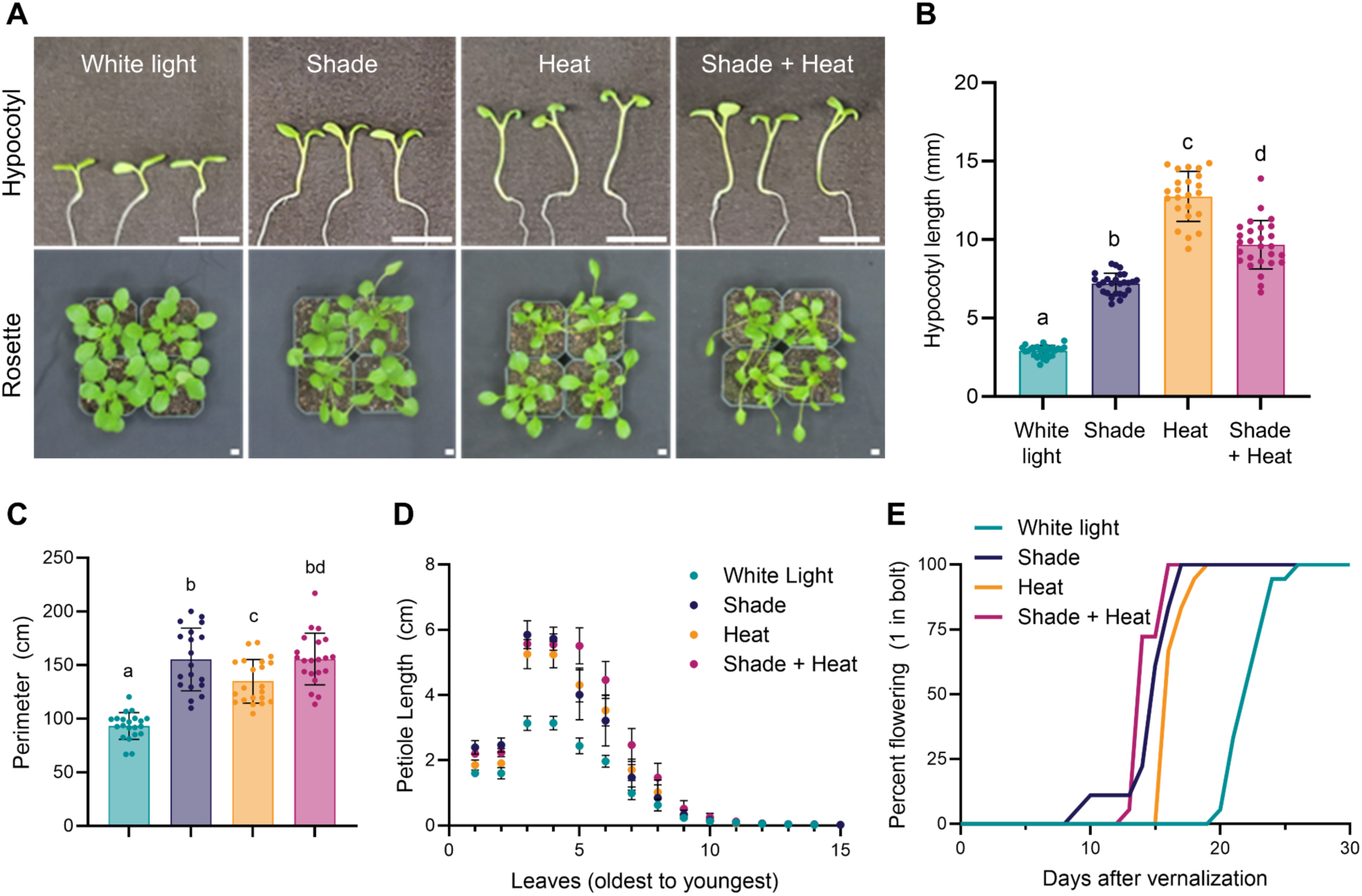
Pennycress increases responses to shade and elevated temperatures. **(A)** Elongation of 5-d-old seedlings grown in white light, shade, heat, and shade + heat. Rosette morphology of plants treated for 6 d with shade, heat, or shade + heat. Scale bar = 1 cm. **(B)** Hypocotyl length of 5-d-old seedlings grown in the indicated treatment for 3 d. Statistical significance determined by 2-tailed Welch’s t-test (± SD, n ≥ 23, p < 0.05). **(C)** Perimeter of 3-week-old rosettes grown in white light, shade, heat, or shade + heat for 14 d. Statistical significance determined by 2-tailed Welch’s t-test (± SD, n ≥ 19, p < 0.05). **(D)** Petiole length of each leaf of a 17-d-old rosette treated for 7 d. Leaves 1 and 2 are cotyledons (n = 14). **(E)** Number of days till flowering. Plants were considered flowering when the bolt ≥ 1 inch (n= 18).

The photomorphogenic response to shade led us to hypothesize that the red-light signaling pathway likely underlies the SAR phenotypes in pennycress and may be initiated by PHYB. Since PHYB co-senses temperature in Arabidopsis, we investigated how pennycress responds to a moderately elevated temperature (28 ℃) and the interaction of shade and temperature ((Burko et al. 2022; Romero-Montepaone et al. 2020; Casal & Fankhauser 2023). Pennycress increased hypocotyl length at 28 ℃, and when in combination with shade (Figure 2). Hypocotyls elongated the most in heat alone, while the combination of shade and heat resulted in an intermediate length between the average length seen in the individual treatments of shade and heat. The response to combined heat and shade contrasts with the additive effect observed in Arabidopsis (Burko et al. 2022), suggesting that pennycress responses to light and temperature are distinct.

Pennycress rosettes were exposed to the various environmental conditions to determine how shade and heat impact other developmental stages. Wild-type pennycress was germinated and grown under control conditions for seven days, then maintained under white light or grown under shade, heat, or shade and heat, for an additional seven days. Shade, heat, and shade and heat-treated plants had increased rosette perimeter (Figure 2) and convex hull area, but lower solidity compared to control conditions (Figure 4), indicating that rosette morphology changes in response to the environment. The changes in rosette morphology are attributed to the increased petiole length of leaves when grown under treatment conditions.

Light quality and temperature are critical environmental cues that influence flowering, and SAR or higher heat can lead to earlier flowering as a competitive or escape response (Wollenberg et al. 2008). Effects of shade and temperature on pennycress flowering time were determined by germinating seeds for one week under white light, vernalizing for three weeks, then moving to white light for four days before starting shade and/or heat treatment for fourteen days. Shade and heat-treated plants bolted earlier than plants grown under control conditions, demonstrating that pennycress alters flowering time in response to environmental cues (Figure 2).

We conclude that pennycress exhibits SAR phenotypes of organ elongation and early flowering, indicating that pennycress is a shade-avoidant species. Together, these experiments support that pennycress alters morphology in response to elevated temperatures and foliar shade, and the aberrant morphology seen at high planting densities may be due, in part, to SAR. Next, we targeted SAR and thermomorphogenesis pathways to determine if we could alter pennycress responses.

### *pif7* has attenuated responses to foliar shade and heat throughout its lifespan

To test if mutating PIF7 could manipulate SAR and thermomorphogenic pathways in pennycress, we sought to use gene editing to disrupt *PIF7* in pennycress. Genomic analysis identified a likely *T. arvense PIF7* homolog (TaPIF7). TaPIF7 has 80% nucleotide similarity and over 90% amino acid conservation compared to *Arabidopsis thaliana* PIF7 (AtPIF7) (Supplemental Figure 3). We generated PIF7 mutants in MN106 with CRISPR-Cas9 targeting the second exon and the bHLH domain (Figure 3). Sequencing the *PIF7* locus identified mutant alleles in transgene-free homozygous mutant lines. *pif7-1* is a deletion of 242 bases that removes 75 amino acids and disrupts the bHLH domain. *pif7-2* is a deletion of 86 bases, which results in a protein that is 122 amino acids instead of 364 amino acids. Due to the severe mutations of *pif7-1* and *pif7-2*, we hypothesize that these are knockout alleles. Importantly, both *pif7* alleles similarly reduced hypocotyl elongation in shade, heat, and shade and heat combined, compared to wild type (Figure 3).

**Figure 3.**
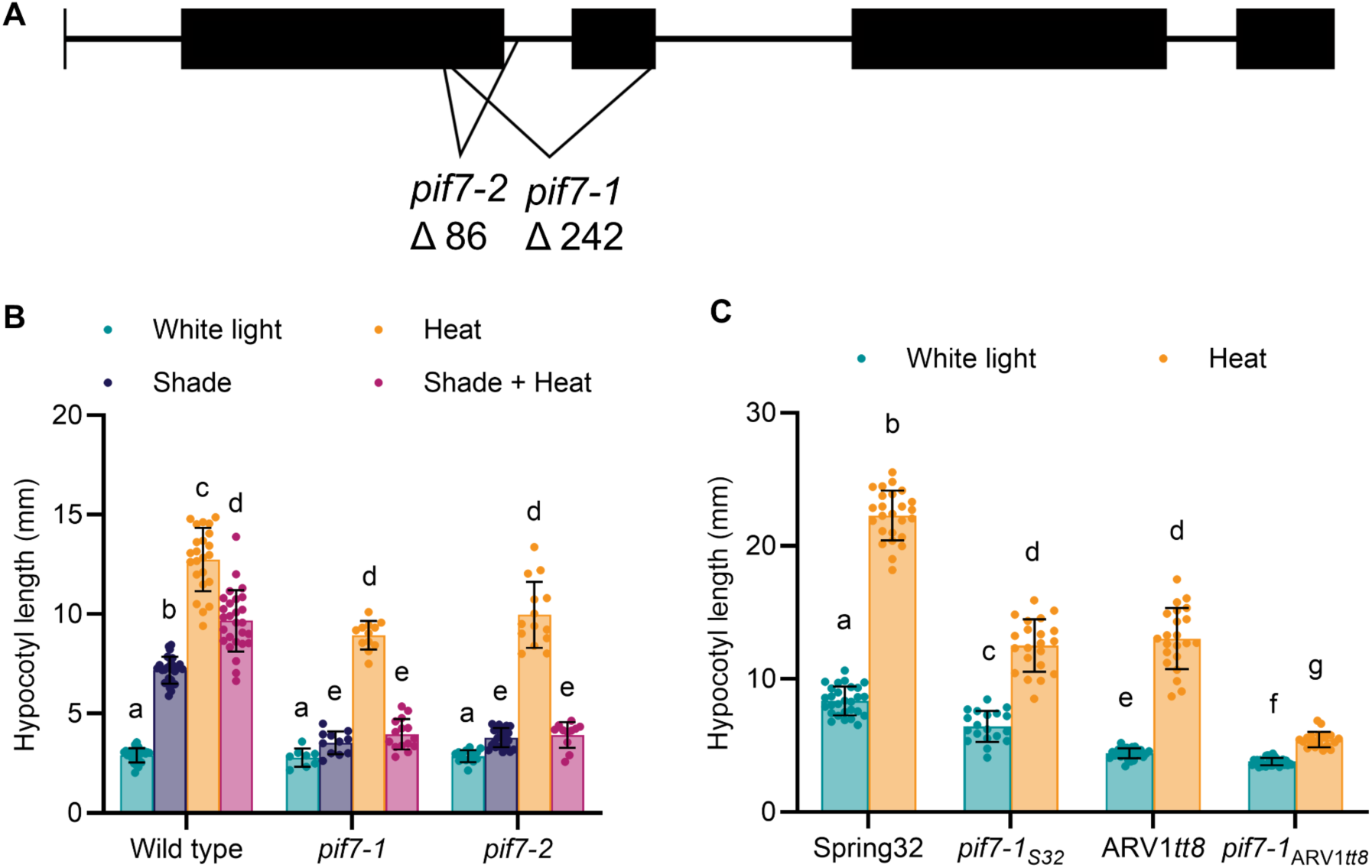
*pif7* has reduced hypocotyl elongation across genetic backgrounds in response to shade and heat. **(A)** PIF7 gene diagram depicting mutations. Exons are black boxes. *pif7-1* and *pif7-2* were generated with CRISPR in the MN106 wild-type background. **(B)** Hypocotyl length of 5-day-old seedlings. Seeds were germinated in white light for 2 d, then grown in shade, heat, or shade + heat, for an additional 3 d. Statistical significance determined by 2-tailed Welch’s t-test (± SD, n ≥ 8, p < 0.05). Data for wild type is replotted from Figure 2B for comparison. **(C)** Hypocotyl length of 5 d-old seedlings, germinated in white light for 2 d, then grown in heat for an additional 3 d. Statistical significance determined by 2-tailed Welch’s t-test (± SD, n ≥ 19, p < 0.05). Common letters indicated no significant difference.

To investigate how loss of PIF7 impacts diverse germplasm, we mutated PIF7 in other pennycress genetic backgrounds. Spring32, a wild-type spring ecotype line, and ARV1*tt8*, an early-generation winter ecotype commercial line, were transformed with the CRISPR-Cas9 constructs targeting PIF7. Strikingly, the *pif7-1* allele was regenerated in both Spring32 and ARV1*tt8*, as well as additional alleles of *pif7* (Figure 3B and Supplemental 3). *pif7-1_S32_* and *pif7-1^ARV1^* mutants reduced hypocotyl elongation in heat in comparison to their parental lines (Figure 3 and Supplemental 3), supporting that the *pif7* phenotype is not unique to the MN106 background and could be useful when introduced into elite breeding lines.

We further characterized the *pif7-1* in the MN106 background to determine if *pif7* had attenuated SAR and thermomorphogenesis across developmental stages, tissues, and flowering time. Rosette morphology was compared between wild type and *pif7-1* in shade, elevated temperature, and combined conditions for 14 days. *pif7-1* was similar to wild-type under control conditions, remaining compact with high solidity, reflecting that *pif7* does not stunt growth in control conditions (Figure 4). When exposed to shade, heat, and the combined conditions, wild-type plants increased rosette convex hull area and reduced solidity (Figure 4). In comparison, *pif7* had reduced convex hull area and increased solidity when compared to wild type under shade, heat, or the combined treatments (Figure 4). Comparison of petiole length of dissected rosettes found that *pif7-1* petioles were shorter compared to wild type under all conditions (Figure 4 and Supplemental Figure 4). The elongation was most prominent in the oldest true leaves as they have been exposed to environmental stress for the longest time. The reduced petiole length of each *pif7* leaf results in an overall more compact plant in shade and elevated temperatures.

**Figure 4.**
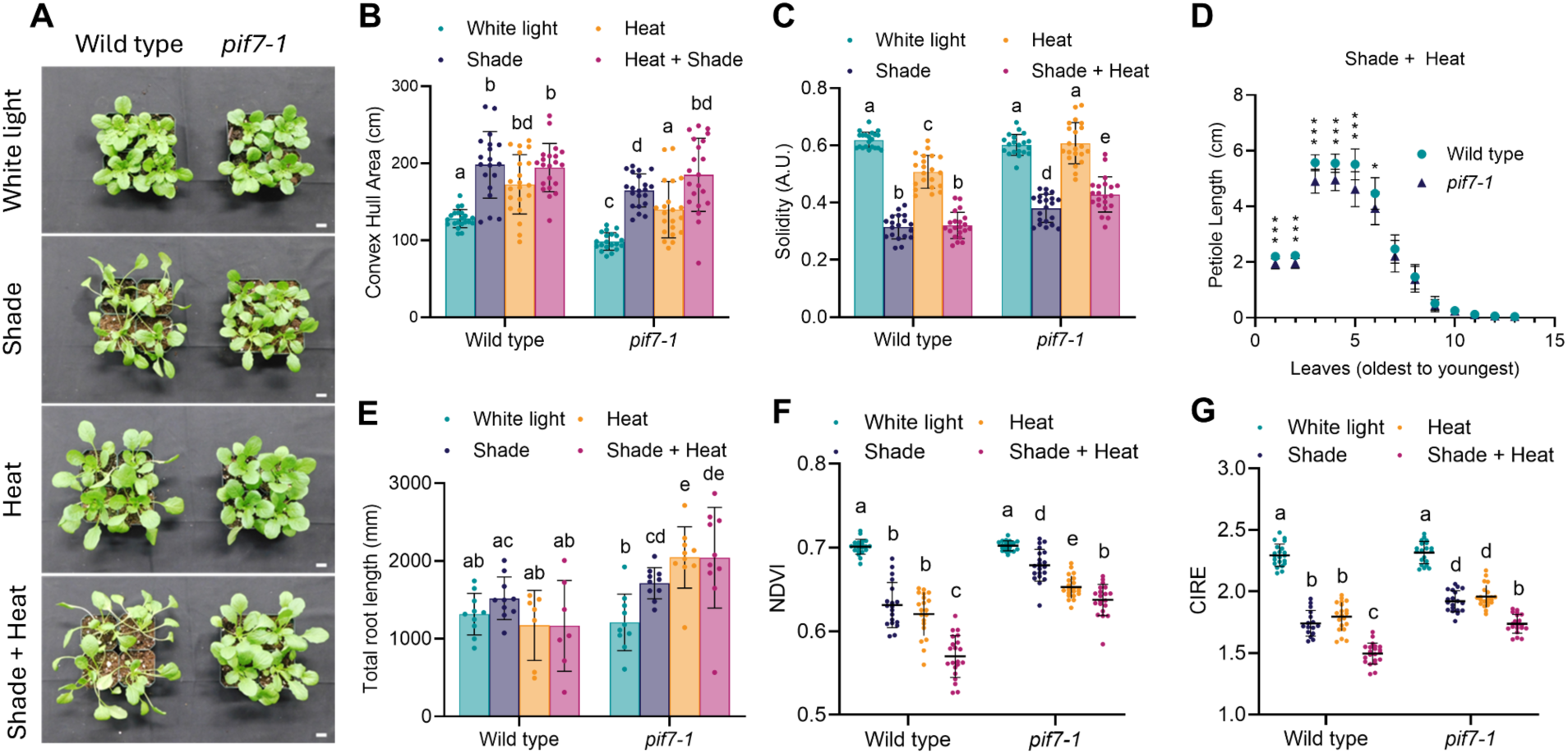
*pif7* attenuates responses to foliar shade and heat throughout vegetative growth. **(A)** Representative plants grown in control conditions for 7 d, then for 14 d in treatment conditions and imaged. **(B)** Rosette solidity of plants grown in control conditions for 7 d, then for 14 d in treatment conditions. Solidity was determined using the shape analysis function from PlantCV. **(C)** The convex hull area of rosettes was calculated using the shape analysis function from PlantCV. **(D)** Petiole length of each leaf of plants grown in control conditions for 10 d, then moved to shade + heat environment for an additional 8 d. Plants were dissected to measure the petiole length of each leaf. Leaves are arranged oldest to youngest; cotyledons are leaves 1 and 2. Statistical significance was determined by 2-tailed Welch’s t-test (± SD, n = 14, p* < 0.05, p** < 0.01, p*** <0.001). **(E)** Total root length of plants grown on MS plates for 7 d, then transferred to turface. Plants were then grown in treatment conditions for 10 d. Roots were washed, scanned, and analyzed with Rhizovision (± SD, n ≥ 7, p < 0.05). **(F)** Seeds were sown in soil and grown in long day control conditions (22 °C, 200 µmol m^-2^ s^-1^, R:Fr =12) for 7 d. On day 7, conditions were changed to shade (22 °C, 200 µmol m^-2^ s^-1^, R:Fr = 0.3), heat (28 °C, 200 µmol m^-2^ s^-1^, R:Fr =12) or shade + heat (28 °C, 200 µmol m^-2^ s^-1^, R:Fr = 0.3). Elevated temperature and the far-red ratio were maintained only during the day. Normalized Difference Vegetation Index (NDVI) and **(G)** chlorophyll index-red edge (CIRE) were measured in dark-adapted plants after 14 d of being grown in the indicated environment. Statistical significance was determined by a 2-tailed Welch’s t-test (±SD, n ≥16, p < 0.05). Common letters indicated no significant difference.

We sought to holistically characterize the impact of mutating *pif7* on plant growth and responses to the environment. Shade and temperature also affect root development, as plants allocate more resources to above-ground growth under these conditions (Salisbury et al. 2007; Rosado et al. 2022; van Gelderen et al. 2018; Gray & Brady 2016). Analysis of the root system found that total root length remained comparable for wild-type plants grown in all conditions, while *pif7-1* had increased total root length in shade, heat, and combined conditions (Figure 4).

Chlorophyll fluorescence indices are associated with physiological health and stress levels (Arief et al. 2023; Ghosh et al. 2023; Murchie & Lawson 2013; Stamford et al. 2023) and were used to estimate the impact of shade, heat, and combined conditions on wild-type and *pif7-1* health. We measured normalized difference vegetation index (NDVI), chlorophyll index-red edge (CIRE), maximum efficiency of photosystem II (F_v_/F_m_), and non-photochemical quenching (NPQ), in wild type and *pif7* grown under shade and heat conditions. Wild type reduced NDVI and CIRE in shade and heat conditions, conveying that these environments may lower chlorophyll content (Figure 4). Both NDVI and CIRE measurements were higher in *pif7-1* than wild type in the same environmental condition, suggesting that *pif7-1* is less sensitive to environmental influences on chlorophyll content (Figure 4). By contrast, the average F_v_/F_m_ of wild type and *pif7-1* in all conditions is within the typical range expected of healthy plants (Supplemental Figure 5) (Murchie & Lawson 2013). Similarly, NPQ increased in shade and heat conditions for wild type and *pif7* mutants, suggesting that plants are activating protective mechanisms to prevent damage from ROS, but is comparable between genotypes (Supplemental Figure 5).

To determine the effects of *pif7* on pennycress morphology, flowering time, and inflorescence architecture, wild type and *pif7-1* were compared under fourteen days of treatment. *pif7-1* bolted earlier under shade, heat, and in combined conditions compared to the control. However, *pif7* was delayed in flowering by 1-2 days compared to wild type in all conditions (Figure 5). At maturity, height was measured to determine if *pif7-1* affected overall plant height. Wild-type plants were taller in shade, heat, and combined conditions than when in control conditions. Mature *pif7-1* plants grown in experimental conditions were also taller than in control conditions, but remained over 10% shorter than wild-type in the comparative condition (Figure 5). This indicates that two weeks of shade and warm temperatures have a lasting impact on plant architecture, and that *pif7* attenuates their physiological impacts.

**Figure 5.**
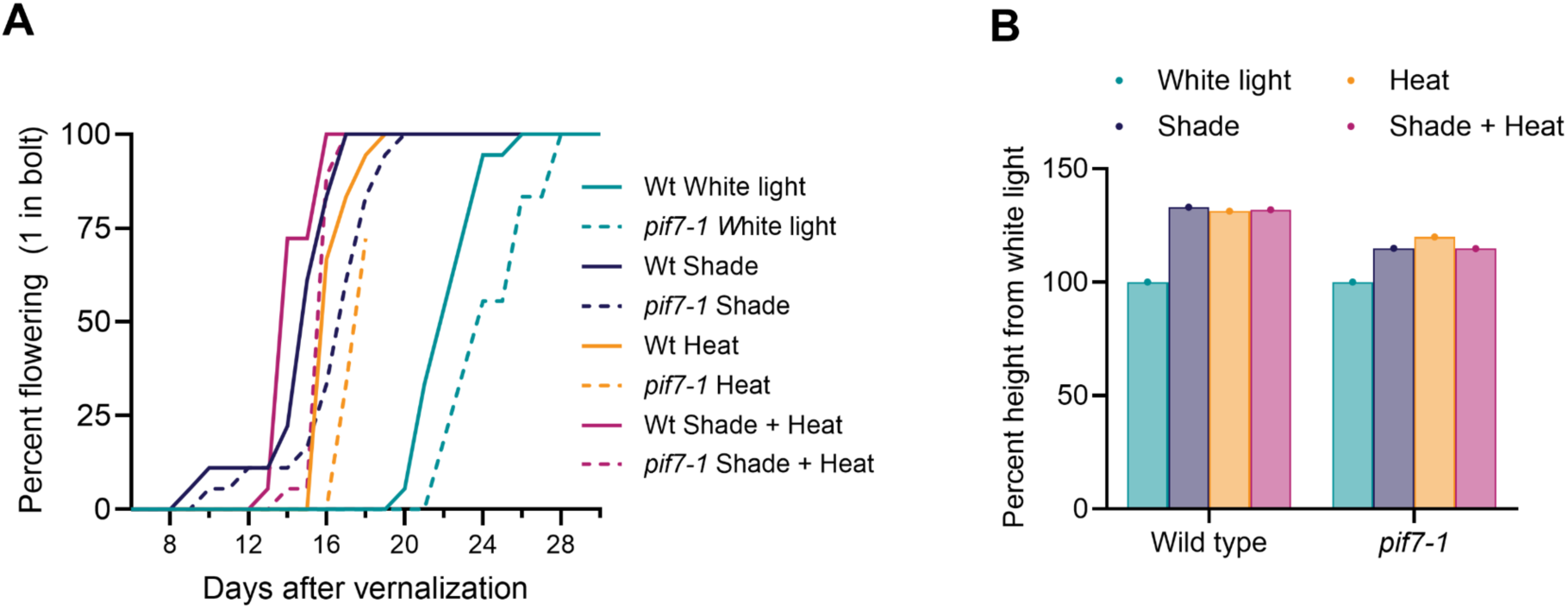
*pif7* attenuates responses to foliar shade and heat through reproductive phases. **(A)** Flowering time of plants grown for 1 week, vernalized for 3 weeks, then moved back to control conditions for 5 d. Plants were then transferred to the indicated treatment condition for 14 d before returning to control conditions. Plants were scored as flowering when bolt height reached one inch (n = 18). **(B)** Plant height was measured at maturity. Height was normalized to plants grown under white light for the duration of the experiment (n ≥ 16).

### *pif7* does not reduce seed oil content, yield, or freezing tolerance

Pennycress is grown for its seed oil content, so we compared seed traits of *pif7-1* to wild-type plants grown in all conditions. *pif7-1* did not alter total yield or thousand seed weight (TGW) of plants when transiently treated for 14 d during vegetative growth (Supplementary Figure 6). Seed oil content on a dry weight basis (DWB) was also unaltered in *pif7-1*. This indicates that mutations in PIF7 do not negatively affect yield or oil content (Supplemental Figure 6).

In Arabidopsis, PIF7 is known to have additional roles in abiotic stress and is a known negative-regulator of DRE-Binding1, a transcription factor important for low-temperature responses (Kidokoro et al. 2009; Lee & Thomashow 2012). PIF7 repression of DREB1 is hypothesized to reduce expression of cold tolerance genes in unstressed conditions and prevent DREB-induced growth reduction. We tested cold and freezing tolerance, as it is crucial for pennycress winter survival as an intermediate crop. *pif7-1* survived freezing challenges similarly to wild type (Supplementary Figure 7), suggesting that loss of PIF7 did not affect cold tolerance up to –10 °C.

### *pif7* has reduced elongation in low light

If intercropped, pennycress will experience low light and shaded conditions as ground level light intensity (fluence rate) and quality (R:Fr) are both reduced in standing corn and dried soybeans (Supplemental Figure 2). In standing corn, light intensity was decreased nearly 10-fold from approximately 1000 µmol m^-2^ s^-1^ down to 100 µmol m^-2^ s^-1^. In standing green corn, the R:Fr was also lowered from 1.4 to 0.6 at ground level. Light conditions in standing corn are various and complex, and are a significant hurdle to overcome during intercropping. We sought to explore any effects of low light intensity on pennycress in a controlled environment.

Elongation of wild type and *pif7* was measured in both seedlings and rosettes when exposed to low light. Hypocotyl elongation in response to a titration of light intensity found that both *pif7* mutants elongated hypocotyls less than wild type in response to reduced light intensity at 20 µmol m^-2^ s^-1^ to 45 µmol m^-2^ s^-1^ (Figure 5). Elongation in low light was also assessed at the rosette stage. Wild type and *pif7-1* were grown under the recommended control light intensity, 300 µmol m^-2^ s^-1^, and low light, 80 µmol m^-2^ s^-1^. Similar to the results for hypocotyl elongation, *pif7-1* elongates petioles less than wild type when grown in low light.

### *pif7* has reduced elongation at high density without impacting yield

To assess if reducing SAR enhances pennycress performance at high densities, *pif7-1* was sown in mini-fields at seeding rates of 5 and 10 lbs per acre and monitored through senescence. The mini-fields were vernalized at 4 *°*C for 3 weeks following a two-week germination and establishment period. After vernalization, the mini-fields were returned to the greenhouse and imaged every five days to monitor ground cover and flowering time. As expected, mini-fields sown at 10 lbs/a had a higher ground cover percentage than 5 lbs/a. All genotypes and planting densities reached around 90% green coverage at 15 d post-vernalization (Figure 7).

**Figure 6.**
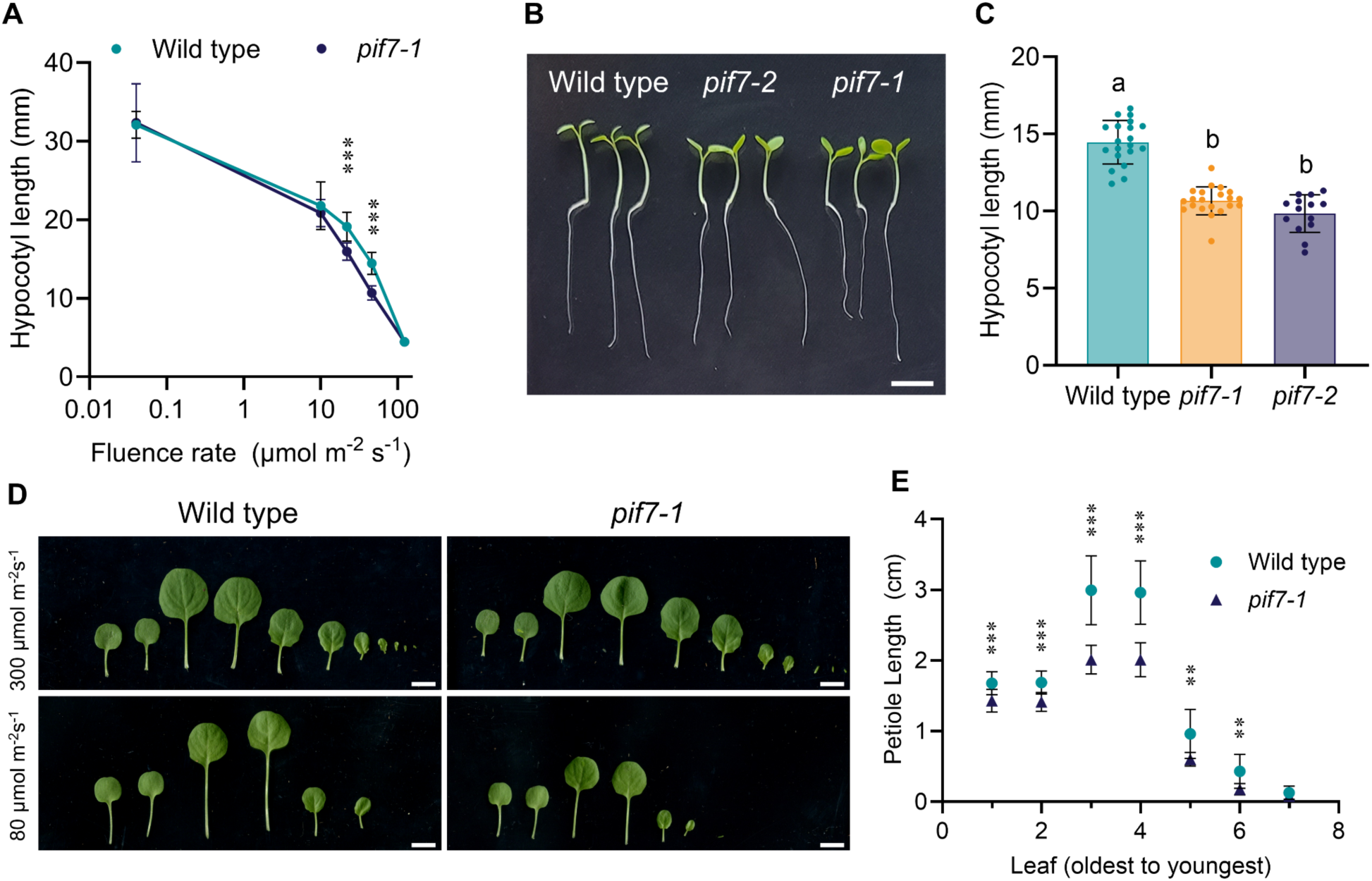
*pif7* has reduced organ elongation in low fluence conditions **(A)** Seedlings were germinated in white light at 120 µmol m^-2^ s^-1^ for 2 d. On day 3, seedlings were grown under various neutral density filters to reduce PAR. Hypocotyls were measured on day 6. Statistical significance was determined by 2-tailed Welch’s t-test. (± SD, n ≥ 14, p* < 0.05, p** < 0.01, p*** <0.001) **(B)** Representative hypocotyls at 45 µmol m^-2^ s^-1^. Scale bar = 5 mm. **(C)** Hypocotyl length of seedlings germinated at 120 µmol m^-2^ s^-1^ for 2 d then grown under 45 µmol m^-2^ s^-1^ for an additional 4 d. Statistical significance was determined by 2-tailed Welch’s t-test (± SD, n ≥ 14, p < 0.05). **(D)** Representative leaves of plants grown under control light and low light conditions. Plants were sown and grown for 1 week at 300 µmol m^-2^ s^-1^. Half of the plants were moved to 80 µmol m^-2^ s^-1^ of light and grown for an additional week, the other half was kept under control light conditions. On day 15, all plants were dissected. Leaves were arranged from oldest to youngest. Scale bar = 1 cm **(E)** Petiole length of each leaf of wild type and *pif7-1* grown at 80 µmol m^-2^ s^-1^. Statistical significance determined by 2-tailed Welch’s t-test (± SD, n ≥ 14, p* < 0.05, p** < 0.01, p*** <0.001). Common letters indicated no significant difference.

**Figure 7.**
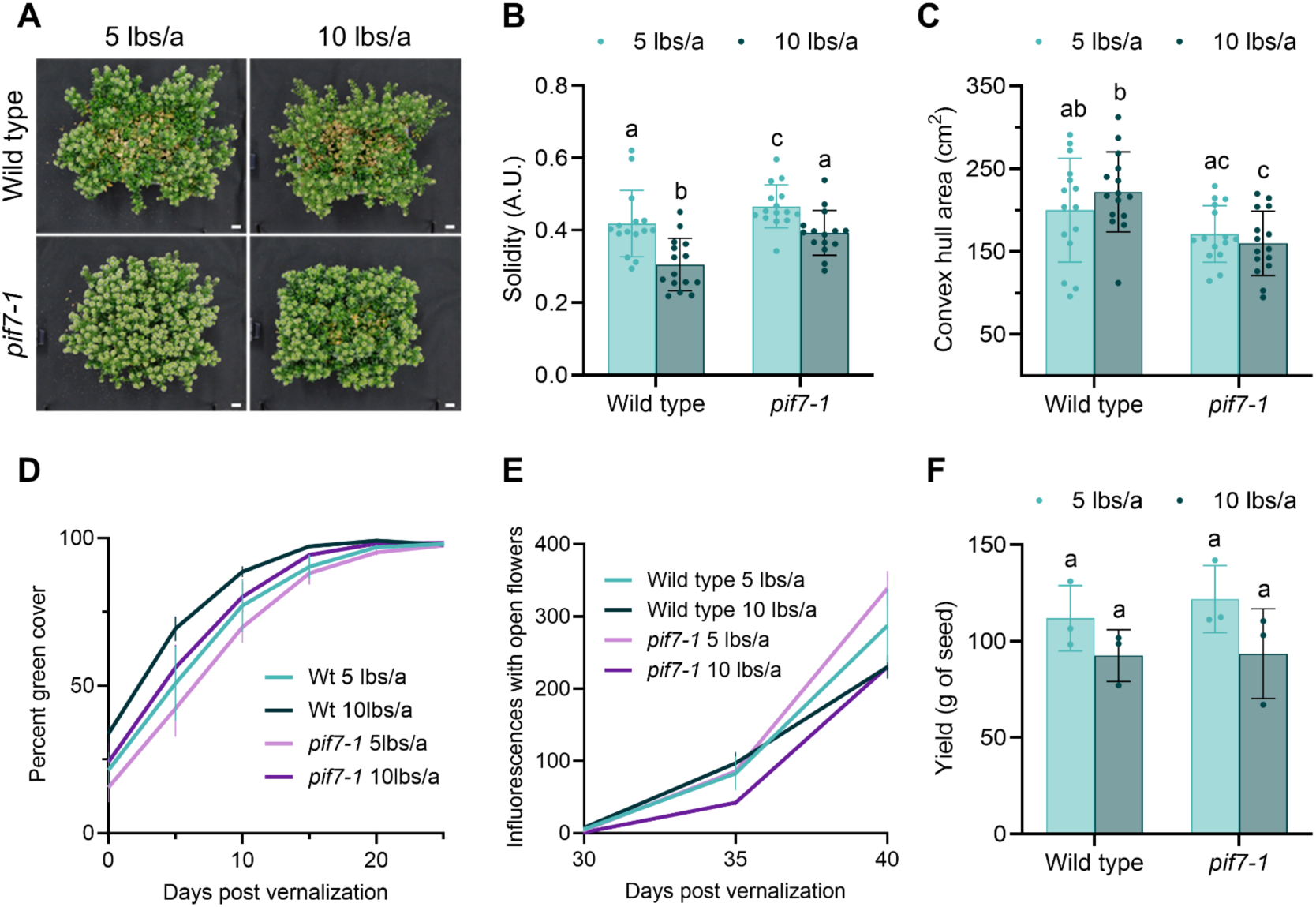
*pif7* attenuates elongation when grown at high density without reducing yield. MN106 and *pif7-1* seeds were sown in 27 L Sterilite containers measuring 59.7 cm x 42.9 cm x 14.9 cm at 5 and 10 lbs/a. The mini fields were grown for 2 weeks, vernalized for 21 d, moved back to standard growing conditions, and imaged every 5 d. **(A)** Mini-fields 40 d after vernalization. **(B)** Five rosettes were pulled randomly from each mini field 11 d after vernalization. Solidity and **(C)** Convex hull area was calculated using the morphology function from PlantCV. Statistical significance determined by 2-tailed Mann-Whitney test (±SD, n = 15, p < 0.05). **(D)** Percent green coverage within the boundary of the mini-field was calculated from overheard images taken every 5 days post-vernalization with PlantCV. **(E)** Inflorescences with open flowers were counted in each mini-field. **(F)** All seeds were harvested for each mini-field to calculate average yield. Statistical significance determined by 2-tailed Mann-Whitney test (±SD, n = 3, p < 0.05). Common letters indicated no significant difference.

Density and genotypic effects on individual plant morphology were assessed at 11 days post-vernalization. This time point was chosen to capture rosette morphology when mini-fields had high green cover, ensuring neighboring plants were touching, but before plants transitioned to flowering. Wild-type rosettes at 10 lbs/a had increased convex hull area and perimeter, but reduced solidity compared to 5 lbs/a (Figure 7). In contrast, *pif7-1* rosettes maintained their rosette shape at both planting densities, as convex hull area, solidity, and perimeter are not statistically different between 5 lbs/a and 10 lbs/a (Figure 7).

All mini-fields had open flowers at 30 d post-vernalization, with wild type at 10 lbs/a averaging the most inflorescences with open flowers. All mini fields reached 100% flowering at 40 days post-vernalization, but varied in the number of inflorescences (Figure 7). *pif7* mini-fields appeared healthier than wild type at both planting densities and flowered uniformly. The center of each wild-type mini-field yellowed and failed to flower at both planting densities; plants along the perimeter were the only ones that flowered, while *pif7-1* had overall less yellowing and clearings. Yield remained statistically the same for wild type and *pif7-1* at all planting densities, showing that reducing SAR responses does not decrease yield.

## Discussion

Here we show that mutating PIF7 reduced SAR and thermomorphogenesis in pennycress with limited impacts on plant health, yield, and seed oil content. Reducing SAR through gene-editing is a rapid and promising strategy to increase crop performance at high-density plantings in diverse environments. *pif7* grown in high-density monocultures elongated less and had more flowers than wild type, demonstrating reduced plasticity to the surrounding environment. This reduced plasticity will increase uniformity of plant growth across millions of acres, even while facing various environmental challenges within different hardiness zones. Increasing planting density has been one of the most effective strategies to increase yield. As leaf angle optimizations have slowed (Elli et al. 2023), planting density improvements will need to be accomplished through alternative mechanisms. Superior vegetative architecture varies greatly between species, and will require unique identification and optimization of genetic influences of each specific trait. Alternatively, reducing the capability of plants to change their architecture in response to high-density plantings is a more broadly applicable strategy. Coupling superior architecture with reduced responses to neighbor competition could be paramount to maximize plant performance at high densities. This study demonstrates the viability of attenuating SAR to control plant architecture and physiological traits.

Pennycress is currently in the early stages of domestication and presents an opportunity to create a variety suited for use across the US agriculture zones. Widespread adoption of pennycress could produce enough material to convert to fuel to meet the 35 billion gallons of sustainable aviation fuel mandate. Pennycress oil has a fatty acid composition well-suited for conversion to biodiesel and biojet fuel, meets the U.S. Renewable Fuels Standard (Fan et al. 2013), and can produce 600 liters ha^-1^ (65 gal ac^-1^) of oil annually without competing for land use with food crops (Moser et al. 2016; Moser 2012). This, combined with ecosystem services provided by growth over the winter when fields typically lay fallow, positions pennycress as a strong candidate crop to meet our future mandated sustainable aviation fuel goals (Fan et al. 2013; Griffiths et al. 2022).

### Pennycress responds to high-density plantings and environmental cues

Here, we have shown that both above- and below-ground biomass was negatively impacted by high-density plantings in the early commercial line ARV1*tt8* and did not increase yield (Figure 1 and Supplemental Figure 1). Further analysis in controlled conditions showed that pennycress is shade and heat-intolerant and responds to simulated canopy shade by elongating organs and flowering earlier. Elongated petioles at the rosette stages increase rosette perimeter and convex hull area, and expose more soil between the leaves (Figures 2, 4). Etiolated plants reduce robust stand establishment and will decrease soil coverage in the field, thereby lowering the potential ecosystem benefits of weed suppression and soil preservation by planting pennycress (Griffiths et al. 2022; Pantazopoulou et al. 2021). The elongation phenotypes in our controlled environments were similar to those in high-density field plots, suggesting that pennycress SAR underlies some physiological alterations seen with increasing density.

Light quality and quantity are both reduced at ground level in fields with a standing crop, which may reduce pennycress stand establishment when intercropped (Supplemental Figure 2). Elongation in response to low light and shade may limit the ability to intercrop current pennycress varieties into existing corn and soy rotations, particularly in northern states where summer crops are often not harvested before September. Delaying pennycress planting until after corn harvest can negatively impact yield (Cubins et al. 2019; Dose et al. 2017; Patel et al. 2021; Bishop & Nelson 2019), but earlier planting can be challenging as temperatures continue to increase during the fall in the Midwest (Us Epa 2021), triggering thermomorphogenesis.

### Altering pennycress morphology by perturbing PHYB signaling

We sought to simultaneously reduce SAR and thermomorphogenesis by manipulating the PHYB signaling pathway and targeted PIF7 as it plays dual roles in elongation to shade and temperature (Burko et al. 2022; Fiorucci et al. 2020; de Wit et al. 2016; Galvāo et al. 2019; Paulišić et al. 2021). CRISPR alleles were generated in multiple backgrounds, including a first-generation commercial line. Interestingly, we generated an identical allele, *pif7-1*, in all three backgrounds. This feat demonstrates that gene editing can generate identical alleles, likely through repair of breaks through microhomology (Schiml et al. 2016).

The wild-type lines (MN106, Spring32, and ARV1*tt8*) all elongated in heat, but to varying degrees. The variability in elongation response suggests that there is natural phenotypic variability to heat, and yet all PIF7 mutants reduced hypocotyl elongation in response to temperature (Figure 3 and Supplemental Figure 3). This evidence also supports the hypothesis that PIF7 is a viable candidate for manipulation across pennycress genetic backgrounds. Further analysis showed that *pif7* alleles in the MN106 background had comparably reduced hypocotyl elongation to shade, heat, and their combination. In summary, our study implies that PIF7 is part of a shade and elevated temperature signaling cascade affecting plant development.

*pif7-1* attenuated petiole elongation, flowering time, and inflorescence stature compared to wild type in all treatment conditions (Figures 4 & 5). Shade and heat alone, or in combination, continued to shorten the flowering time in *pif7-1* compared to wild type in control conditions (Figure 5). The slight impact on flowering is critical to ensure compatibility of this crop with the planting of summer cash crops. Pennycress holistically responds to the environmental cues by altering root growth, similar to Arabidopsis (Rosado et al. 2022; van Gelderen et al. 2018). Diversion of critical resources to shoot growth is hypothesized to enhance competitiveness in suboptimal light quality. Interestingly, *pif7* continues to increase total root length in shade and elevated temperature, suggesting it has dampened the competitive growth reprogramming response to shoots while increasing root length (Figure 4). Greater root length enhances plant resource capture of water and nutrients by increasing soil exploration and root-soil contact, thereby improving plant resilience and nutrient catch as a cover crop (York et al. 2013; Griffiths et al. 2022).

### Pennycress health and yield is not strongly impacted by shade and temperature

Elongation is a strategy to outgrow shade-casting neighbors or to protect organs from hot soil. We found that *pif7* or reduced SAR does not negatively impact yield and plant health. This study sought to understand if elongation provided a physiological benefit to plants in our test environments by measuring chlorophyll indexes commonly used to assess plant stress (Murchie & Lawson 2013; Maxwell & Johnson 2000; Arief et al. 2023). Wild-type and *pif7-1* individual plants had similar F_v_/F_m_ measurements near the idealized 0.8 measurement in control conditions, shade, heat, and shade plus heat, indicating that these conditions are not stressful enough to damage PSII (Supplemental Figure 5). NPQ is a protective mechanism that increases when plants experience high light or other abiotic stresses (Ghosh et al. 2023; Murchie & Lawson 2013; Stamford et al. 2023). NPQ increased similarly in the test conditions between wild type and *pif7-1*, suggesting that these plants are activating protective mechanisms to prevent damage from ROS, and this response is preserved in *pif7* (Supplemental Figure 5). Similarly, seed quantity, weight, and oil content were not significantly altered in *pif7* compared to wild type in control and experimental conditions (Supplemental Figure 6). These data support the idea that *pif7* does not reduce yield.

Overall, we did not find evidence of growth or photosynthetic impairment in *pif7*, as *pif7-1* remained comparable to wild type when grown in control conditions. We believe *pif7* does not have a significant negative effect outside of suppressing SAR and heat responses, and therefore this mutation is viable for integration into elite breeding lines.

### The potential for *pif7* to improve interseeding

Low light intensity and shade are critical challenges for pennycress establishment in intercropping scenarios (Supplementary Figure 2). When interseeded into standing corn, pennycress will face the complex environment of low fluence and low R:Fr (Supplementary Figure 2). The etiolated morphology that pennycress adopts when grown in standing corn is hypothesized to contribute to poor establishment and reduced yield (Supplemental Figure 2). Reducing elongation in low light and shade may improve interseeding success. Our study demonstrates that, in controlled low fluence environments, pennycress phenocopies morphology in reduced R:Fr by similarly elongating hypocotyls and petioles (Figure 5). *pif7* had reduced hypocotyl and petiole elongation in low light compared to wild type and was not allele specific. Our data is the first to implicate *pif7* function in low fluence and suggests that canonical red light signaling has a role in growth responses to low fluence. This novel finding implicates the PHYB-PIF signaling hub as a regulator of morphology responses to low light.

### Potential to increase planting density

Our ultimate goal is to reduce the undesirable morphogenic changes associated with SAR, allowing plants to maintain ideal architecture when grown at high density. We tested *pif7* performance in a common garden experiment by sowing mini-fields at 5 lbs/a and 10 lbs/a, both densities higher than the recommended 3 lbs/a, in a temperature-controlled greenhouse. Our mini-field experiments also showed that *pif7* had reduced rosette elongation at 10 lbs/a and is statistically the same as wild type at 5 lbs/a in rosette morphological characteristics (Figure 7). This shows that SAR and light quality contribute to the elongated phenotypes seen at high-density plantings. The *pif7* mini-fields looked healthier than wild type due to yellowing and flowered uniformly in the mini-fields. Importantly, yield was not decreased by mutating *pif7,* supporting a hypothesis that we can improve vegetative growth responses to the environment without a yield drag.

### Summary

We have used the foundational knowledge from the Arabidopsis red-light signaling pathway to reduce plasticity to environmental cues in pennycress. Creating a PIF7 mutant resulted in attenuated elongation in shade and elevated temperatures, and under low light intensities associated with interseeding. This study supports that reducing SAR is a promising strategy to optimize plants for planting under challenging conditions found during intereseeding and at higher densities.

## Methods

### Plant materials

MN106 is the primary wild-type line used throughout the paper, and Spring32 and ARV1*tt8* are additional germplasm used to generate additional PIF7 mutants. ARV1*tt8* is an early commercial line with a mutation in TRANSPARENT TESTA 8 (Koirala et al. 2023). *pif7-1* is a novel allele generated in MN106, Spring32, and ARV1*tt8* backgrounds. *pif7-2* was an additional allele generated in the MN106 background, while *pif7-3* was generated in Spring32 background and *pif7-4* was generated in the ARV1*tt8* background. All alleles were developed using CRISPR gene editing as described below. MN106 is the parental background unless otherwise denoted with a subscript.

### Field trials

The field trial was conducted at Western Illinois University in Macomb, IL, USA (40.4885, -90.6902). The preceding crop was Cereal rye, which was harvested on June 26th, 2023. 50lb. N (28% UAN)+ 10lbs. Sulfur (Thiosulfate) was applied as a liquid on September 26. Treflan was applied on all plots at a rate of 1 qt./acre and double incorporated with tillage. Seed was prepared into planting packets. Pennycress seed was broadcast on September 29th, 2023, with ½ inch of water applied within 24 hours of planting. Plots were 4’ x 10’ in size and sowed at 3, 5, 7, and 9 lbs/acre.

Five rosettes were removed from each plot 64 days after planting on December 04, 2023, after plants slowed their growth. Rosettes were imaged with an overhead camera to analyze rosette morphology.

On May 15, 2024, at maturity, five whole plants from the center of each plot, including the root system, were harvested. All plants were individually imaged. The number of tillers arising from the basal rosette was manually counted. Plants from the same plot were dried and weighed as a group to calculate biomass.

One soil core was taken from the middle of each plot and directly over one plant, ensuring that the cored section represented the whole plot. Soil cores were taken from the field on March 27th, 2024, 179 days post-planting using a bulb and garden planter with 6” depth x 3” diameter dimensions. Cores were placed in plastic bags and stored at -20 ℃ until washing. After defrosting in a water bath, the soil cores were carefully washed by hand, and the roots cleaned and separated from particulate organic matter. Washed roots were then stored in clean plastic bags at 4 ℃ and were imaged in the following days using an Epson Expression 12000XL flatbed scanner equipped with a transparency unit (Epson America Inc, Los Alamitos, CA, USA). Roots from the soil core were carefully separated in a water-filled plastic tray into individual plants by the tap root. Roots were placed on square plates (150 mm × 150 mm) filled with a 10 mm layer of deionized water with one tap root per tray. Unconnected roots from neighboring plants that had no tap root were placed in a separate square plate. All roots were spread out using forceps to avoid overlap between roots. Six trays were imaged simultaneously with the scanner using the Epson Scan 2 software (Epson America Inc, Los Alamitos, CA, USA) at a resolution of 600 dpi as a JPG file with 99% quality. The individual root images were cropped out of the main image, and root measures were computed using RhizoVision Explorer v2.0.2 (Seethepalli et al. 2021). In the software, the tap root length, lateral root length, and tertiary root length were classified by root diameter threshold ranges of > 0.4 mm, 0.4 - 2 mm, and < 2 mm, respectively. The image thresholding level was set to 200, and the root pruning threshold was set to 20 pixels. After scanning, the root material was placed in a paper envelope and dried at 60 ℃. In this way, root length density for the plot was determined after aggregating individual plant data together by core. Root length per plant was determined by averaging the root measures by number of tap roots per plot.

### Drone Imaging and Analysis

Drone images were taken after the hard frost period throughout the spring to harvest starting on February 27, 2024, and finishing on May 27, 2024. The whole field was imaged every 7 to 10 days with 12 flights. The drone used for data collection was a DJI Mavic 3M equipped with a multispectral sensor camera. Flights were pre-programmed using DJI Pilot 2 and were conducted at 17 m height above the ground with an image resolution of 0.75 cm/pixel. Images were collected at an approximate flight speed of 1.5 m/s. There was 70% side overlap and 80% frontal overlap in the collected images. The time of the day for each flight was around 9 am CDT, ensuring the weather conditions were similar. Images were assembled into orthomosaics using WebODM v. 2.5.

Image analysis was conducted on the orthomosaic images using a custom Open-CV Python code (Griffiths 2025). A reference scale was established for each orthomosaic using the known field distance, and a scaling factor was applied to ensure consistent dimensions across plots. Then the images were rotated, ensuring each image had the same orientation. Individual plots were then cropped out as a grid layout, and plot names were assigned to the images. To differentiate between bare soil and vegetation, a soil mask was generated using the Blue-Green Index (BGI). Pixels with BGI < 0.15 were classified as soil. Additionally, a green leaf index (GLI) was used to differentiate green and senesced vegetation based on a GLI threshold of 0.2 (>= 0.2 green, < 0.2 senesced). The indices used are defined as:

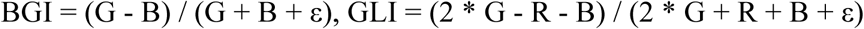

Where R, G, and B are the red, green, and blue channels of the image, respectively, and ε is a small constant to avoid division by zero. The percentages of green tissue and senesced tissue were calculated relative to the total pixel count within the plot. To ensure accurate cropping and thresholding, a reference image was generated for each plot by overlaying the indices on the original orthomosaic for manual validation.

### Seed sterilization and growth conditions

Seeds were sterilized with a 30% bleach and 0.1% Triton solution for 15 minutes and washed three times with sterile water. For bulk sterilization, seeds were gas sterilized by arranging seeds in a single layer and exposed to chlorine gas for 5 hrs. Seeds were plated on ½ strength Murashige and Skoog (MS) medium and stratified at 4 °C for 3-5 d. Plates were grown at 130 µmol/m-^2^/s^-1^ in 12:12 light:dark cycles at 22 °C for 4-7 d then vernalized at 4 °C with 12:12 light:dark for 3 weeks, unless otherwise noted.

Hypocotyl measurements were taken by growing seeds for 2 d at 22 °C 12:12 light:dark control conditions, then exposed to the experimental conditions for the indicated number of days. White light conditions were a 12 hr day/ night at a constant 22 °C. For heat treatment, plates were grown at 28 °C during the day with a 12 hr day/night cycle and 22 °C night temperature. Canopy shade was simulated by adding a far-red light during the daytime to reduce the R:Fr to 0.3. A constant temperature of 22 °C was maintained at plant level by reducing the chamber settings to 20 °C during the day to compensate for the heat given off from the far red lights. For the shade + heat treatment, plates were incubated under far red light during the day at an R:Fr of 0.3. A daytime temperature of 28 °C was maintained by adjusting the chamber to 25 °C during the day when the supplemental far red lights were on and the night temperature was 22 °C. Temperature readings were collected with a HOBO meter to ensure chambers were at the proper temperature.

Plants grown for rosette measurements were sown directly into soil in 3.5 in pots. Light intensity was 200 µmol/m-^2^/s^-1^ with a 16:8 light/dark cycle at 22 °C. Plants had to have visible true leaves before beginning experimental conditions. A daytime temperature of 28 °C was the heat treatment. Shade was simulated with the addition of far red to reduce the R:Fr to 0.3. The chamber temperature was set to accommodate the heat generated by the far red lights, which was 22 °C (shade only) or 28 °C (shade + heat), and night temperatures were 22 °C. Rosettes were imaged on the indicated day.

### Plant transformation and selection

Plants were transformed using *Agrobacterium tumefaciens* strain GV3101 by submerging inflorescences in an Agro solution, then vacuum infiltrated under 27 inches of mercury for at least 10 minutes (McGinn et al. 2019). Inflorescences were wrapped with cling wrap and placed in the dark overnight. In the morning, plants were returned to standard growth conditions and allowed to mature.

T_1_ seeds were selected by growing on 50µg/mL of hygromycin for 2 weeks. Resistant plants were vernalized and rescued to soil. Plants were genotyped to confirm the presence of CRISPR and mutations to the target gene.

### Generating CRISPR alleles

Two guide RNAs were designed with Cas-designer (Park et al. 2015) to target the PIF7 coding region. Guide RNAs (gRNA) were cloned into a pHEE401 CRISPR-containing vector as described. The first gRNA with PAM sequence is TCTTGGGCGTCTTTCGAATCTGG. The second gRNA plus PAM sequence is CGATTCACAACGAGTCTGAAAGG. The pHEE401 containing gRNAs and CRISPR was transformed into pennycress lines: MN106, Spring 32, and ARV1*tt8*. T_1_s were grown for 14 days on ½ MS agar plates containing 50µg/mL hygromycin for selection. T_1_s were genotyped using primers to amplify the region spanning both gRNA targets and by sequencing the PCR products. Lines containing desirable mutations were propagated until homozygous for the mutation and the absence of transgenic elements. Tiling PCR was used to confirm the complete absence of CRISPR-CAS9 in the final lines.

### Image analysis

Whole seedlings were laid out on an MS agar plate with a ruler in the same plane as the plants. The plate was backlit with a lightboard and imaged with an overhead-mounted camera. Hypocotyls were individually measured using ImageJ.

Rosettes were photographed with an overhead-mounted camera on the indicated day. A color card was included in every image to serve as a size reference and allow for color correction if necessary. Images were processed with PlantCV (v 4.2.1) by white balancing each image and then thresholding to isolate the plant from the background (Gehan et al. 2017). The analyze.size function was used to measure shape characteristics and size. The chip size of the color card was measured using the transform.detect_color_card function. The chip size in pixels was used to convert appropriate results to metric measurements.

Mini-fields were imaged with an overhead-mounted camera with a color card in every image. Machine learning was used to train the naive.bayes function of PlantCV to distinguish between the plant and background. Only plant and background material within the perimeter of the mini-field box was used to calculate percent ground cover.

### Density trials

Holes were drilled into Sterilite Holes were drilled into Sterilite 14.9 cm H for drainage, and each is considered a mini-field. Three independent mini-fields of each planting density and genotype were arranged at random in the greenhouse. Seeds were broadcast, allowing for random spacing of seeds. The mini-fields were grown for 2 weeks to allow seeds to germinate before vernalizing for 3 weeks. The mini-fields were fertilized once a week in the greenhouse.

Fields were imaged every 5 days after vernalization. Inflorescences were counted using the cell-counter ImageJ plugin to score open and unopened inflorescences. After senescence, the entire mini-field was harvested to calculate total yield per mini-field.

### Chlorophyll Fluorescence measurements

Seeds were sown and grown in long day conditions for 7 days, then changed to the appropriate treatment condition and grown for an additional 14 days. Rosettes were imaged with the PhenoVation CropReporter imaging system. Plants were dark-adapted for a minimum of 20 minutes before imaging. NPQ, NDVI, and color analysis were taken. Each indice was quantified using the corresponding function in PlantCV.

### Freezing trial

MN106 and *pif7-1* were grown under controlled conditions in a growth chamber maintained at 22°C with a 16-hour light/8-hour dark photoperiod. Seedlings were grown for two weeks at 22°C before being transferred to a vernalization chamber set at 4°C with an 8-hour light/16-hour dark photoperiod for three weeks to induce cold acclimation. The experiment was repeated across five independent batches, each consisting of 12 seedlings per genotype, resulting in a total of 60 seedlings per line for survival evaluation.

Following cold acclimation, seedlings were transferred to a programmable environmental chamber (LU-113, Espec Corp., Hudsonville, MI, USA) for the freezing treatment. Plants were initially held at 4°C for 30 minutes, followed by a gradual temperature reduction at a rate of 2°C per hour until reaching -3°C. This temperature was maintained for 16 hours to promote ice nucleation and pre-freezing conditioning. Subsequently, the temperature was further decreased to -10°C at the same rate (2°C per hour), and plants were held at this target freezing temperature for 2 hours. After freezing, the temperature was gradually increased back to 4°C at 2°C per hour, where plants were held for 1 hour before being returned to standard growth conditions (22°C, 16-hour light/8-hour dark) for recovery.

Plant survival was assessed 1–2 weeks post-treatment, based on visible regrowth and meristem viability. Plants were considered “survived” if new leaf growth emerged and the meristem remained viable. This experimental design enabled a comparative assessment of freezing tolerance between MN106 wild-type and *pif7* mutant lines under controlled freezing conditions.

### Seed oil content

Total seed oil content and moisture were determined by nondestructive TD-NMR on 450 mg samples of whole pennycress seed using a Bruker MQ40 and Minispec Plus version 7.0.0 software to calculate weight percent oil and moisture. TD-NMR was performed on the samples and standards with a Bruker Minispec MQ40 with a 39.95 MHz NMR frequency and 40°C magnet temperature. The 90° and 180° pulse lengths were 11.48 µs and 22.98 µs, respectively. A 23° detection angle, gain of 44 dB, pulse attenuation of 15 dB, recycle delay of 2 s, window 1 of 0.055 ms, window 2 (τε) of 7 ms, and 724 magnetic field steps were set to employ a Hahn spin echo effect for signal collection. Each sample employed 16 scans at a receiver gain of 59. Weight percent oil and moisture calibration curves were constructed within the Bruker software for oil standards (20% - 40% oil) and moisture levels (6% - 11% moisture). For sample analysis, 450 mg of whole pennycress seed was placed into 10 mm flat-bottom NMR tubes, tempered at 40°C in a heat block for 30 minutes, and scanned in the MQ40. The Minispec Plus software applied the standard curves, and the results were exported to Excel. The percent oil on a dry weight basis was calculated by dividing the milligrams of oil in each sample by the dry weight of each sample and multiplying by 100.

## Author contributions

Vanessica Jawahir and Dmitri Nusinow designed this project. Vanessica Jawahir wrote the manuscript, designed, performed, and analyzed experiments. Salma Adam performed and analyzed experiments. Marcus Griffiths collected and analyzed soil cores, and drone images and root morphology in field and controlled experiments. Tad Wesley and Win Phippen designed and performed the in-field density experiment and collected drone images throughout the field season. Wesley and Phippen assisted with all field sample collections and analyzed seed oil content. Nicholas Heller provided the field conditions to measure light. Bhabesh Borphukan and Karen Sanguinet designed and performed the freezing tolerance experiment. All authors edited and reviewed the manuscript.

---

**Supplemental Figure 1.**
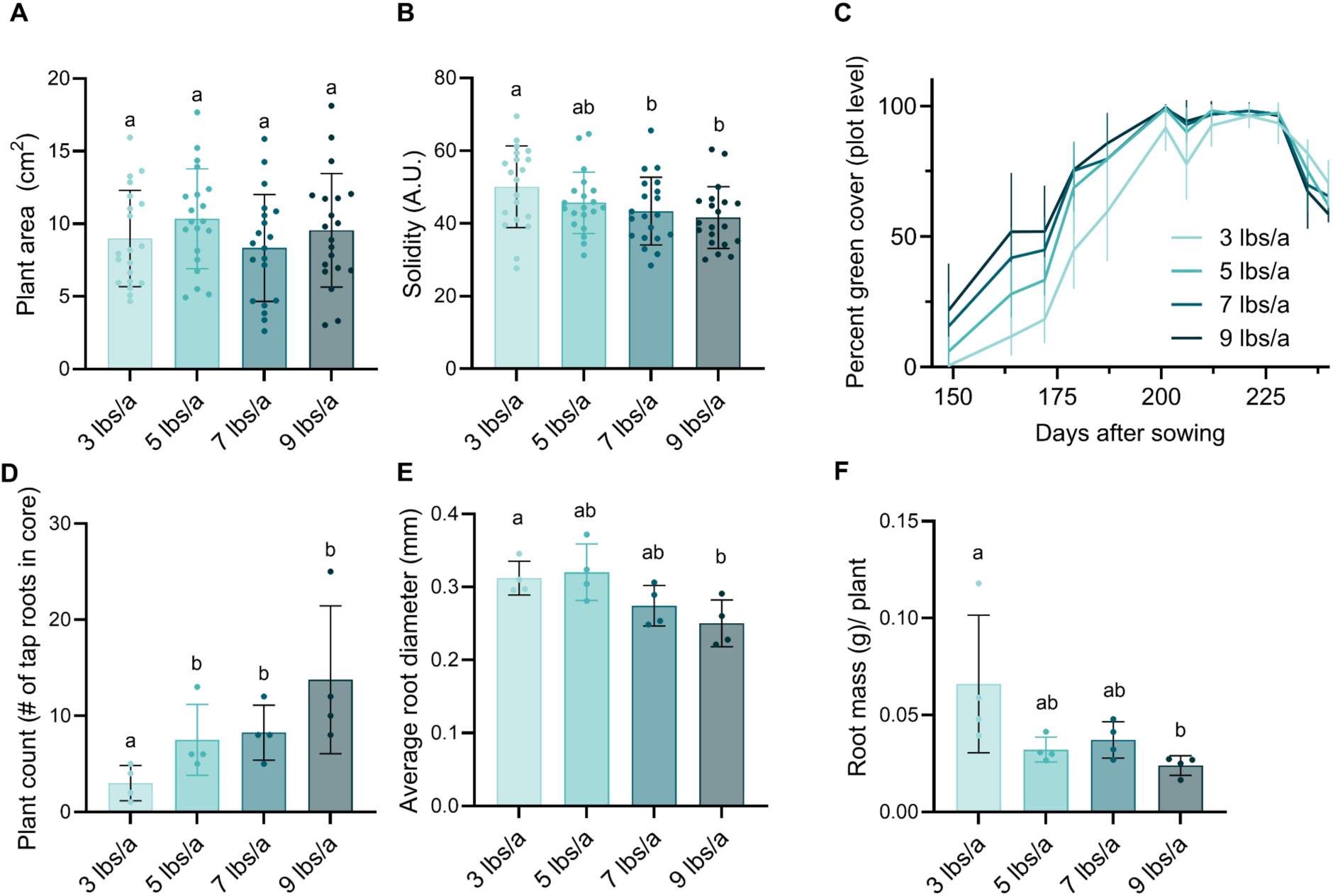
Additional field measurements. Five individuals were removed from each density plot and imaged on 12-04-2023, 64 days after planting. **(A)** Plant area and **(B)** solidity were calculated using the shape analysis function from PlantCV. Statistical significance was determined using a 2-tailed Mann-Whitney test (±SD, n = 16, p < 0.05). **(C)** The percent green cover of each plot was calculated using drone images to estimate plant coverage of the soil. **(D)** Soil cores were taken from the center of each plot. Tap roots from each core were counted to measure the number of plants per core. Statistical significance was determined using a 2-tailed Wilcoxon rank sum test (±SD, n = 4, p < 0.05). **(E)** Root traits were measured using Rhizovision. The root diameter of all roots in the core was averaged. Statistical significance was determined using a 2-tailed Wilcoxon rank sum test (±SD, n = 4, p < 0.05). **(F)** Roots were washed and tried to measure biomass. Total biomass was divided by the number of tap roots in the core to estimate root mass per plant. Statistical significance was determined using a 2-tailed Wilcoxon rank sum test (±SD, n = 4, p < 0.05).

**Supplemental Figure 2.**
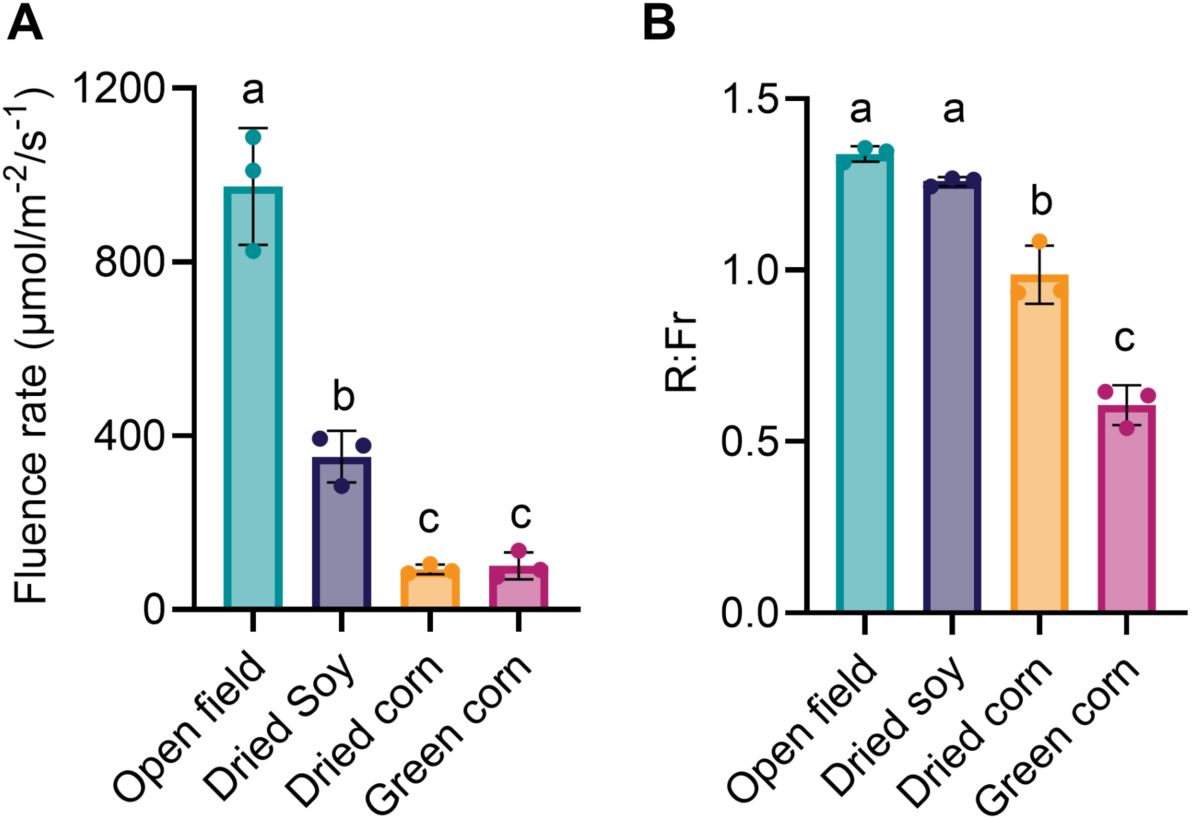
Light intensity and quality are altered in interseeded fields and affect yield. **(A)** Light intensity and **(B)** quality were measured on a partly cloudy day on 10-23-2023 between 1-3 pm with an Li-180. Standing corn measured approximately 9 ft in height. Light readings were taken by walking down a straight line and taking 10-12 measurements approximately every 10 feet. These measurements were averaged for each repetition, n=3. Statistical significance was determined using 2-tailed Welch’s t-test (±SD, n = 3, p < 0.05). Common letters indicated no significant difference

**Supplemental Figure 3.**
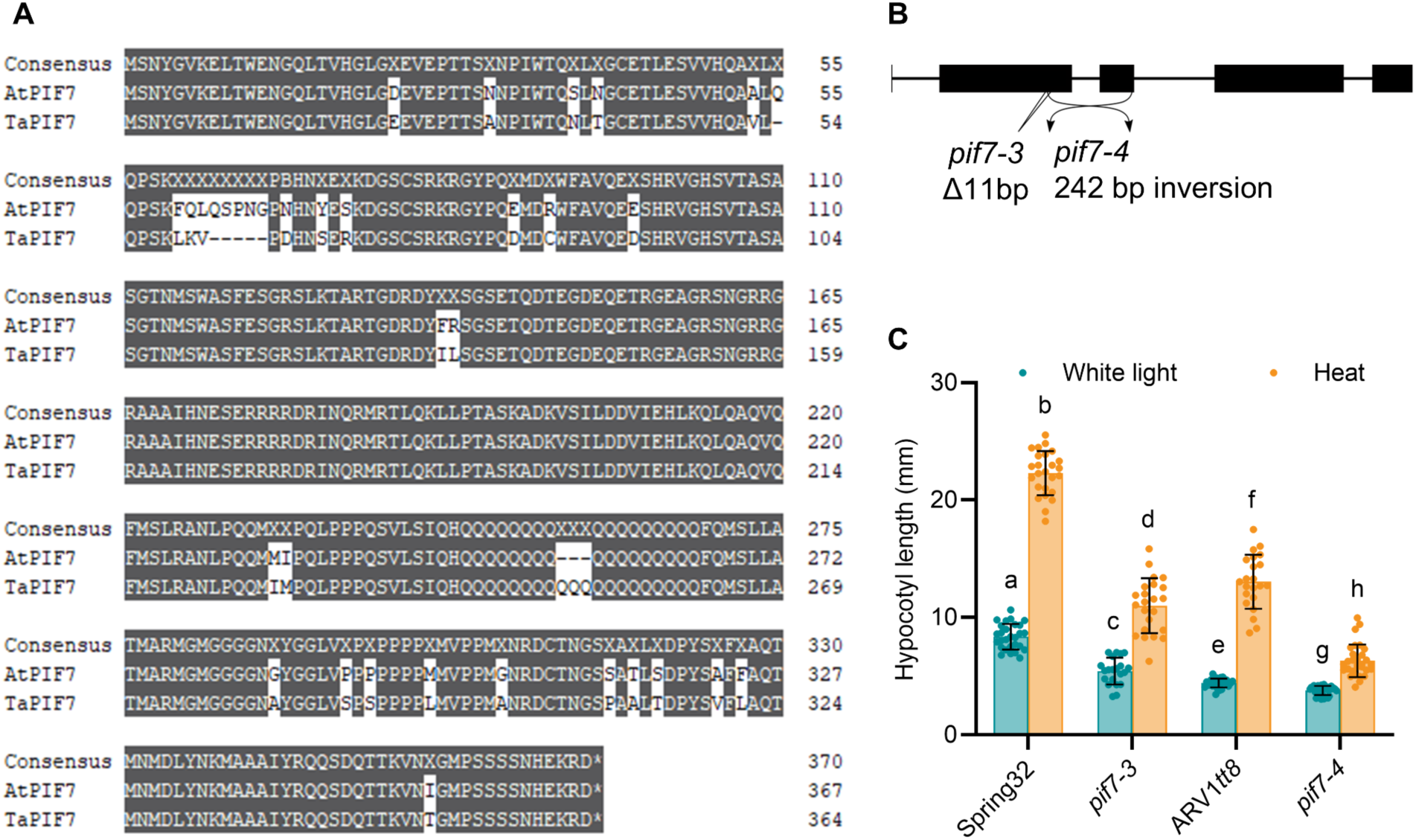
**(A)***Thlaspi arvense* PIF7 (TaPIF7) is highly similar to *Arabidopsis thaliana* PIF7 (AtPIF7), and *pif7* has reduced elongation in heat. **(A)** Amino acid alignment of pennycress PIF7 (TaPIF7) and Arabidopsis PIF7 (AtPIF7). Agreements with the consensus are highlighted. **(B)** PIF7 gene diagram depicting additional alleles generated by CRISPR-CAS9. Exons are black boxes. *pif7-3,* in the Spring32 genetic background, is a small deletion of 11 bases that results in a frameshift. *pif7-4*, in the ARV1*tt8* background, is an inversion of the gene sequence between the gRNA targets. **(C)** Hypocotyl length of 5 d-old seedlings, germinated in white light for 2 d, then grown in heat for an additional 3 d. Statistical significance determined by 2-tailed Welch’s t-test (± SD, n ≥ 21, p < 0.05). Common letters indicated no significant difference.

**Supplemental Figure 4.**
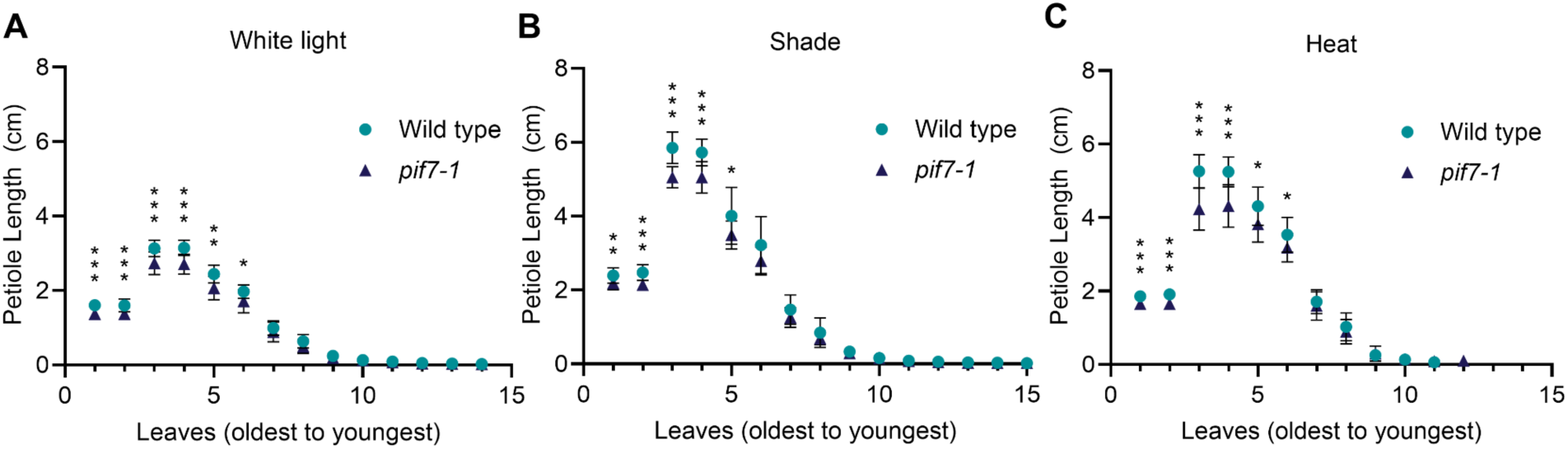
p*i*f7*-1* has reduced responses to environmental conditions. Petiole length was measured for each leaf of a rosette grown in **(A)** white light, **(B)** shade, and **(C)** heat. Plants were initially grown in control conditions for 10 d before moving to treatment conditions for 8 d before dissection. Leaves are arranged oldest to youngest; cotyledons are leaves 1 and 2. Statistical significance was determined by 2-tailed Welch’s t-test (± SD, n = 14, p* < 0.05, p** < 0.01, p*** <0.001).

**Supplemental Figure 5.**
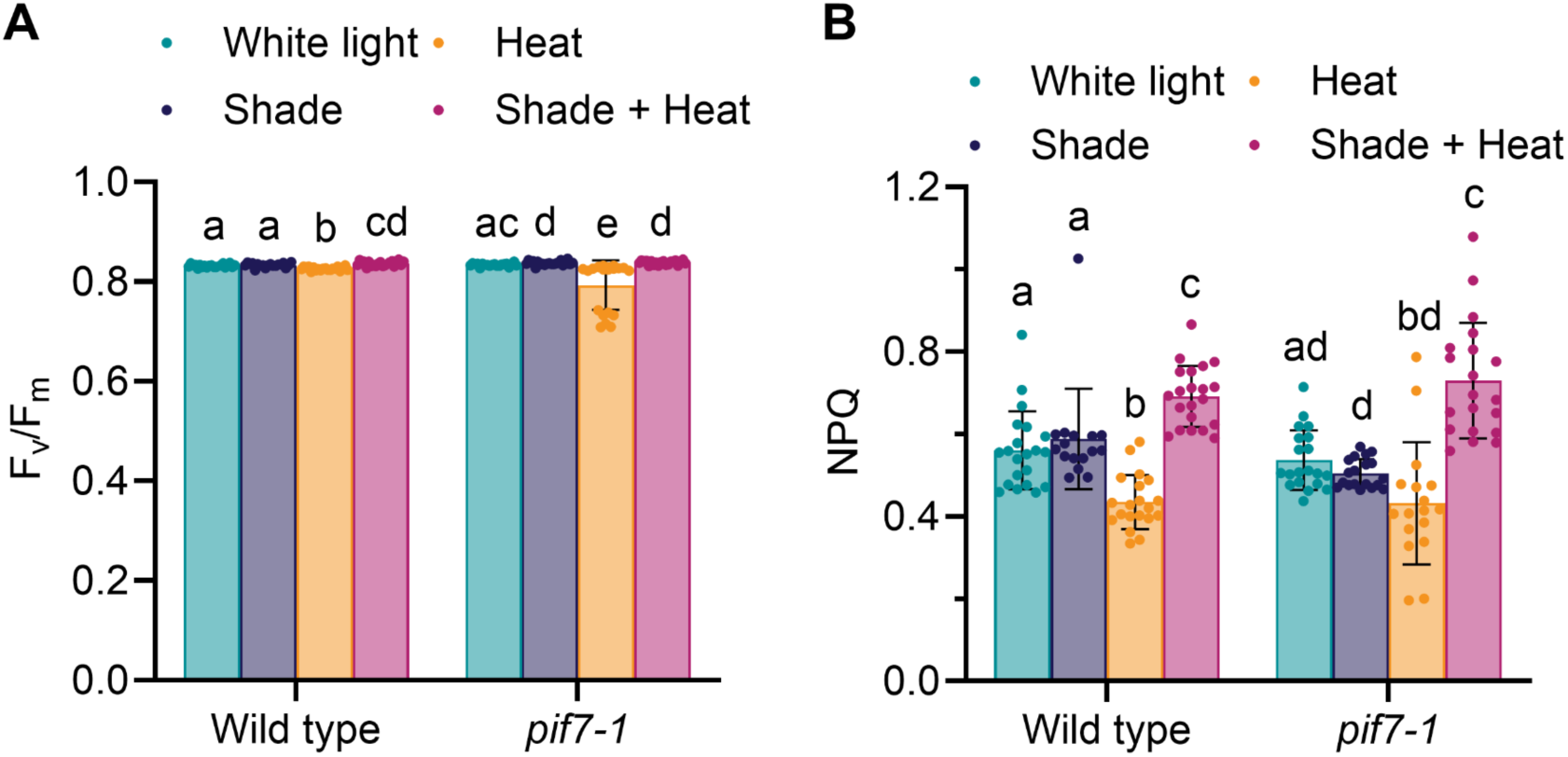
Chlorophyll fluorescence measurements of plants grown in shade and heat. **(A)** F_v_/F_m_ and **(B)** NPQ were measured in dark adapted plants after 14 d of being grown in the indicated environment. Seeds were sown in soil and grown in long day control conditions (22 °C, 200 µmol m^-2^ s^-1^, R:Fr =12) for 7 d. On day 7, conditions were changed to shade (22 °C, 200 µmol m^-2^ s^-1^, R:Fr = 0.3), heat (28 °C, 200 µmol m^-2^ s^-1^, R:Fr =12) or shade + heat (28 °C, 200 µmol m^-2^ s^-1^, R:Fr = 0.3). Elevated temperature and the far-red ratio were maintained only during the day. Statistical significance was determined by a 2-tailed Welch’s t-test (±SD, n ≥16, p < 0.05). Common letters indicated no significant difference.

**Supplemental Figure 6.**
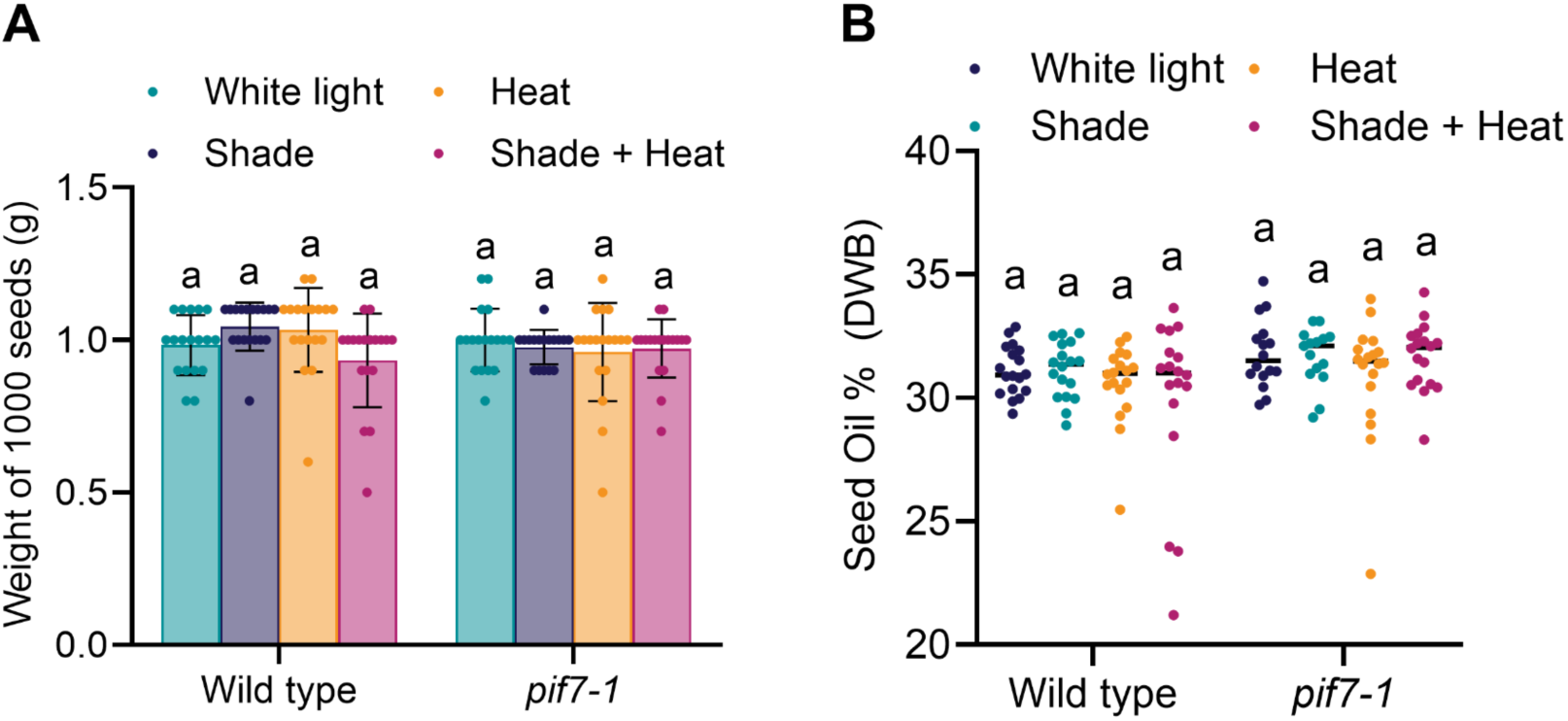
p*i*f7 does not alter seed weight and oil content. Plants at the rosette state were transferred to the indicated condition for 14 d after vernalization, then moved back to control conditions to mature before seeds were harvested and measured. (A) One thousand seeds from each plant were weighed. Statistical significance was determined by 2-tailed Mann-Whitney test (± SD, n ≥ 17, p < 0.05). (B) Seed oil percentage on a dry weight basis (DWB) was measured in seeds. Statistical significance was determined by 2-tailed Mann-Whitney test (± SD, n ≥ 15, p < 0.05).

**Supplemental Figure 7.**
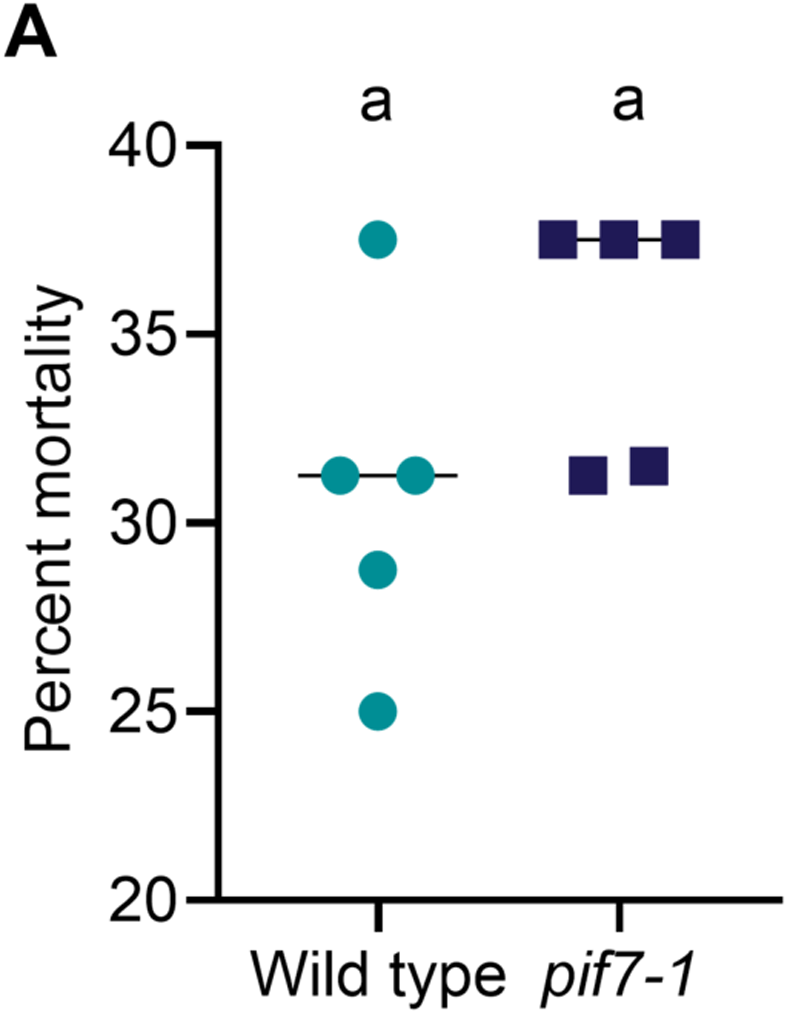
p*i*f7 maintains freezing tolerance. Plants were grown for 2 weeks, vernalized for 3 weeks at 4 °C, then subjected to freezing treatment down to -10 °C. Plants were returned to the greenhouse and evaluated for survival after a 10 d recovery period. 16 plants of each genotype were tested for freezing tolerance in 5 independent tests. Statistical significance was determined by a 2-tailed Mann-Whitney test (± SD, n ≥ 5, p < 0.05). Common letters indicated no significant difference.

